# EASI-PASS: An accessible pipeline for linking functional imaging and mRNA profiling

**DOI:** 10.64898/2026.08.21.746328

**Authors:** Jonnathan Singh Alvarado, Crystian Massengill, Josh Stern, Oren Amsalem, Bettina Ventura, Ahram Jang, Sarah Cook, Ana Veliche, Praneel Sunkavalli, Diti Patel, Juliana Colaccino, Kathryn Evans, Yuhan Wang, Mark L. Andermann

**Affiliations:** Division of Endocrinology, Metabolism, and Diabetes, Beth Israel Deaconess Medical Center, Boston, MA, 02115; Department of Neurobiology, Harvard Medical School, Boston, MA, 02115; Program in Neuroscience, Harvard Medical School, Boston, MA, 02115; Department of Pathology, Boston Children’s Hospital and Harvard Medical School, Boston, MA, 02115; Division of Nutritional Sciences, Cornell University, Ithaca, NY, 14853

**Author notes:** Correspondence (M.L.A.). Janelia Research Campus, HHMI, Ashburn, VA, 20147. Contributed equally to this study (Crystian Massengill, Josh Stern, Oren Amsalem).

## Abstract

We developed EASI-PASS, a reliable, high-throughput method for estimating the molecular identity of functionally characterized cells by merging live imaging with subsequent fixed-tissue imaging using conventional microscopes. Our method matches the shapes and locations of thousands of densely imaged cells between large (>1 mm^2^) functional images and a thick, expanded, and cleared EASI-FISH tissue volume to assess gene expression. This approach is more efficient than alignment to thin sections and recovers the molecular identity of ∼78% of cells. In acute brain slice imaging from the mouse parabrachial nucleus during optogenetic stimulation of long-range spinal inputs, we observed fine-scale specificity in the molecular identity of spinorecipient neurons. In the awake mouse visual cortex, we observed distinct arousal modulation and spatial falloff in correlations within and across interneuron classes. Thus, EASI-PASS provides reliable and efficient alignment of cellular activity with molecular identity.

## Introduction

Advances in optical recordings allow neuroscientists to track the activity of the same thousands of cells over days to weeks *in vivo*^1–8^, while advances in spatial transcriptomics allow molecular profiling of thousands of cells in similar-sized chunks of brain tissue^9–13^. However, multimodal measurements combining functional imaging with spatial mRNA profiling in the same cells have not become mainstream due to several technical challenges. For example, functional images and post-hoc *in situ* hybridization images differ due to complex tissue deformation during slicing and handling, and differences in each cell’s fluorescence intensity and spatial profile across modalities. Strategies to address these problems have recently emerged^14–18^ but typically require specialized hardware and can be challenging to implement in-house. Further, limited matching efficiency and a focus on smaller images (typically <0.5 mm^2^) have resulted in relatively low numbers of matched cells per experiment^15,17–20^.

To enable reliable, large-scale and efficient cellular-level matching between functional imaging and mRNA measurements, we developed EASI-PASS (**EASI**-FISH^21,22^ **P**ost-hoc **A**lignment by **S**patial **S**egmentation). Our approach should be feasible for many labs to implement, as it provides entry points for live imaging in both slice and *in vivo* preparations and is compatible with conventional two-photon (2P) microscopes for functional recordings and standard confocal microscopes for post-hoc tissue imaging. Several factors contribute to the effectiveness of our method. First, EASI-PASS cell registration is based on the location and shape of each segmented cell within an image rather than on computationally intensive pixel-by-pixel alignment of images. Second, EASI-PASS is tolerant to minor tilt mismatches, as it aligns functional images to expanded and cleared thick tissue slabs (up to 500 µm thickness, using the EASI-FISH^21^ protocol). Third, imaging of the same sparse fluorescent markers in live tissue and after post-hoc EASI-FISH provides effective spatial anchors for initial coarse alignment. Finally, the use of the soma-restricted calcium indicator Ribo-GCaMP8s^23,24^, while not required, improves cell matching efficiency due to the similarity in cell shape to the *ex vivo* cytoDAPI stain used for EASI-FISH segmentation^21^. Ribo-GCaMP8s protein can also be cleared more effectively than GCaMP6 or GCaMP8, freeing an extra color channel for mRNA imaging. In addition to cross-modal alignment, our pipeline includes modules for stitching, segmentation, multi-round FISH registration (adapted from the original EASI-FISH workflow to better handle standard confocal imaging volumes), and extraction of molecular signals.

The basic EASI-PASS workflow can be summarized as follows: (1) provide standard image files (TIFF) from functional recordings and post-hoc EASI-FISH volumes; (2) place corresponding landmarks at manually matched anchor points such as shared sparse markers in each dataset (BigWarp^25^); (3) segment cells in the functional and *ex vivo* datasets and use the cell masks to align and match cells; and (4) register *in situ* hybridization rounds, extract gene expression values, and merge all information into a single output table for further analysis.

EASI-PASS yielded new biological insights across two very different proof-of-principle applications. EASI-PASS effectively matched cells between acute brainstem slice calcium imaging and EASI-FISH (72% of 20,397 imaged cells across 7 mice). Here, we identified molecular markers enriched in parabrachial nucleus neurons driven by optogenetic stimulation of spinal afferents. In a second application, EASI-PASS matched cells between *in vivo* cortical calcium imaging in the visual cortex of awake mice and EASI-FISH (83% of 27,152 imaged cells across 5 mice). In particular, we simultaneously tracked the activity of ∼600 molecularly defined interneurons as well as thousands of other neurons per mouse, revealing diverse arousal coupling and spatial scales of correlated activity within and across interneuron classes. Together, this work establishes EASI-PASS as a relatively straightforward approach for linking activity dynamics with gene expression of thousands of cells using conventional 2P and confocal microscopes.

## Results

### EASI-PASS pipeline structure

We designed EASI-PASS to integrate preprocessing, data handling, and alignment of the functional and *ex vivo* imaging into one workflow (**Figure 1A**). We describe the pipeline for the case where 2P calcium imaging is used to record the activity of cells across multiple depths and then these cells are aligned to a single thick EASI-FISH slab of cleared, expanded tissue. The pipeline is organized into three arms (**Figure 1B**). In the functional arm (green), the user provides a 2P image of one of the imaged planes (additional planes can be registered automatically). In the *ex vivo* arm (purple), the user provides an image volume for each round of hybridization chain reaction^26–28^ (HCR) mRNA labeling (typically 5 color channels per round across 1-10 rounds). Cell masks are extracted^29,30^ from 2P and EASI-FISH datasets using soma-targeted Ribo-GCaMP8s^23,24^ and cytosolic DAPI (cytoDAPI^21^), respectively. Manually placed landmarks for putatively matched cells in each image provide an initial coarse alignment. From here, the cross-modal arm (orange) improves alignment in global and local phases by optimizing mask overlap. Final decisions on cell matches to the first round of mRNA labeling are made using an overlap metric or a metric based on neighbor relations (Soma-print^16^). In parallel, each subsequent round of mRNA labeling is aligned to the first round using a modified version of the EASI-FISH registration workflow. All processed cells, mRNA values, and match quality scores are then combined into a single feature table for downstream analyses. Below, we describe this pipeline in greater detail for *in vivo* recordings and then illustrate its use in two example applications in brain slices and *in vivo*.

**Figure 1.**
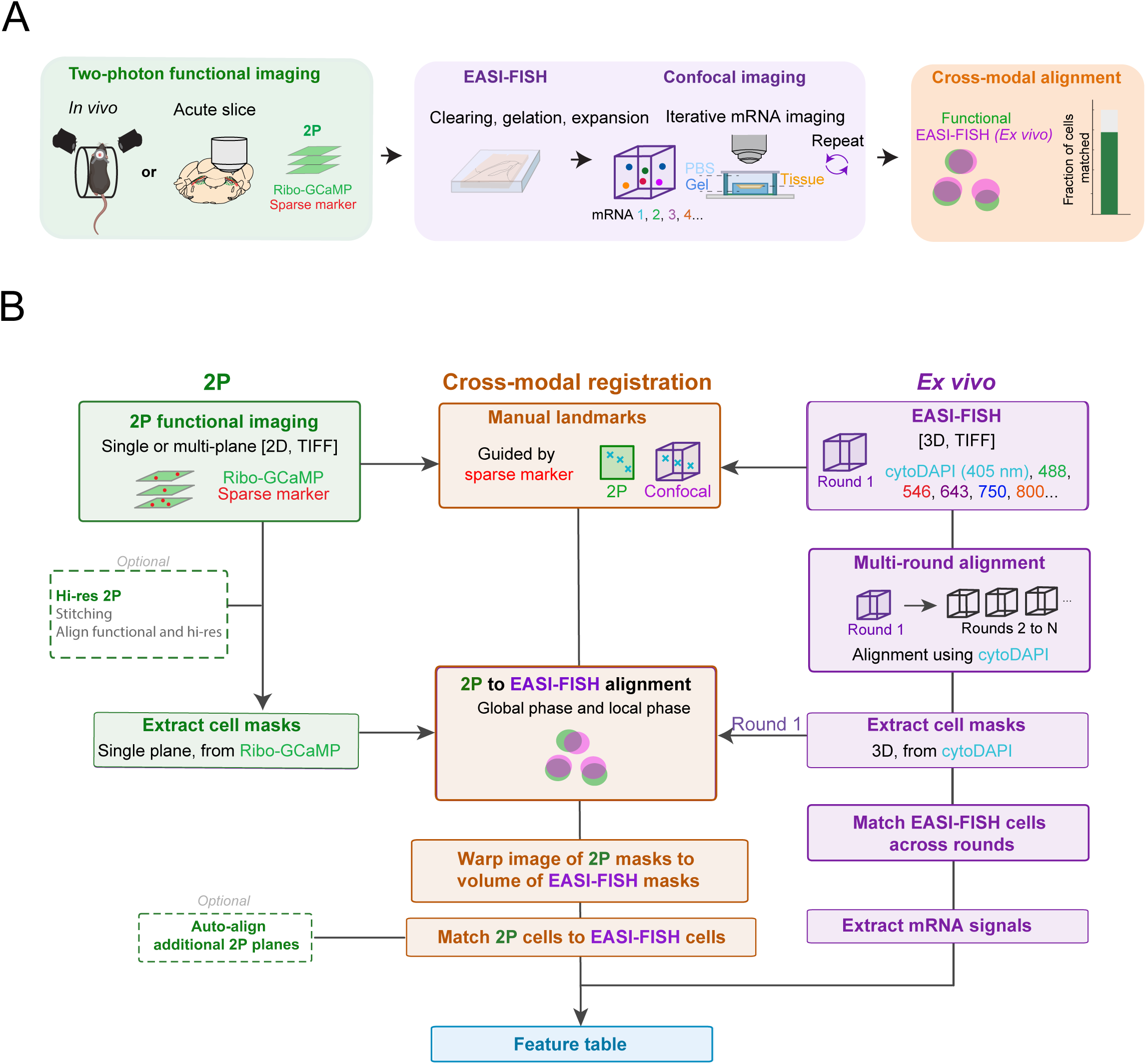
EASI-PASS pipeline overview. **(A)** Two-photon (2P) functional images (green) are registered to *ex vivo* EASI-FISH volumes (purple) by aligning their cell masks (orange). **(B)** EASI-PASS workflow. Cell masks are extracted from 2P imaging planes and EASI-FISH confocal volumes. Cross-modal alignment begins by placing manual landmarks at locations of the same sparse markers in both datasets, then proceeds through registration based on mask overlap, followed by cell matching. Extracted mRNA values, alignment metrics, and match calls are combined into a feature table.

### Cross-modal registration approach

We first illustrate the EASI-PASS pipeline for aligning *in vivo* 2P imaging planes to post-hoc EASI-FISH volumes. To image neurons densely *in vivo* together with a sparse marker for alignment, we co-injected AAV viruses to express Ribo-GCaMP8s in all neurons and tdTomato in *Sst-* expressing neurons in the visual cortex of Sst-Cre mice. We then implanted a cranial window above visual cortex to allow for multi-plane *in vivo* two-photon imaging in layer 2/3. After the final imaging session, we perfused the mouse and extracted a tissue block (3 × 3 × 0.25 mm, from 20-270 μm below the cortical surface) cut tangential to the imaged region, using the cranial window imprint as the physical reference for the cutting angle (**Supplemental Figure 1A** and see Methods). Because EASI-PASS matches cells within a thick, expanded slab rather than to thin sections, this block only needed to be approximately tangential. We trimmed the block to ∼2 × 2 × 0.25 mm and ran the first round of the EASI-FISH protocol.

The EASI-FISH protocol readily removed Ribo-GCaMP8s fluorescence, while tdTomato fluorescence remained intact (**Supplemental Figure 1B-C**) and was clearly visible in both the 2P and *ex vivo* EASI-FISH images, providing a sparse marker for cell alignment. Corresponding landmark pairs were placed manually in BigWarp^25^ to allow for initial coarse alignment. Subsequently, the alignment was refined through a global affine transformation followed by stages of increasingly local warping of each subregion of the image (**Figure 2A**). Each stage began by identifying masks putatively belonging to the same cells in both the 2P images and the EASI-FISH volume. We extracted a square patch containing each cell and nearby cells in either image, and computed an overlap metric (intersection over union, IoU, see **Figure 2B**) between the 2P Ribo-GCaMP8s masks and corresponding EASI-FISH cytoDAPI masks, over a range of rigid 3D shifts. The patch size and range of shifts varied with the size of the subregion being aligned. Well-matched cells showed a clear IoU peak at a specific shift, and these high-confidence matches were used to compute an affine transformation. This strategy was repeated at progressively finer scales, with a maximal level of warping allowed at each scale to avoid overfitting (**Figure 2C, see Methods**). For each iteration in this process, additional local shifts were accepted if a sufficient number of cells had matches and if match overlap improved within a given subregion. These accepted shifts were then interpolated across tiles to form a smooth displacement field. Across stages, the displacement fields grew more structured as the imaging plane deformed from a flat sheet into a surface with increasingly smaller-scale corrections (**Figure 2D**). Several independent quality metrics showed clear improvements across iterations (**Supplemental Figure 2A-D**) as 2P and EASI-FISH masks came into mutual alignment across the field of view (**Figure 2E-F**). The same workflow, applied to 2P imaging in acute brain slices containing the parabrachial nucleus (PBN), produced similar results (**Supplemental Figure 2E-F**).

**Figure 2.**
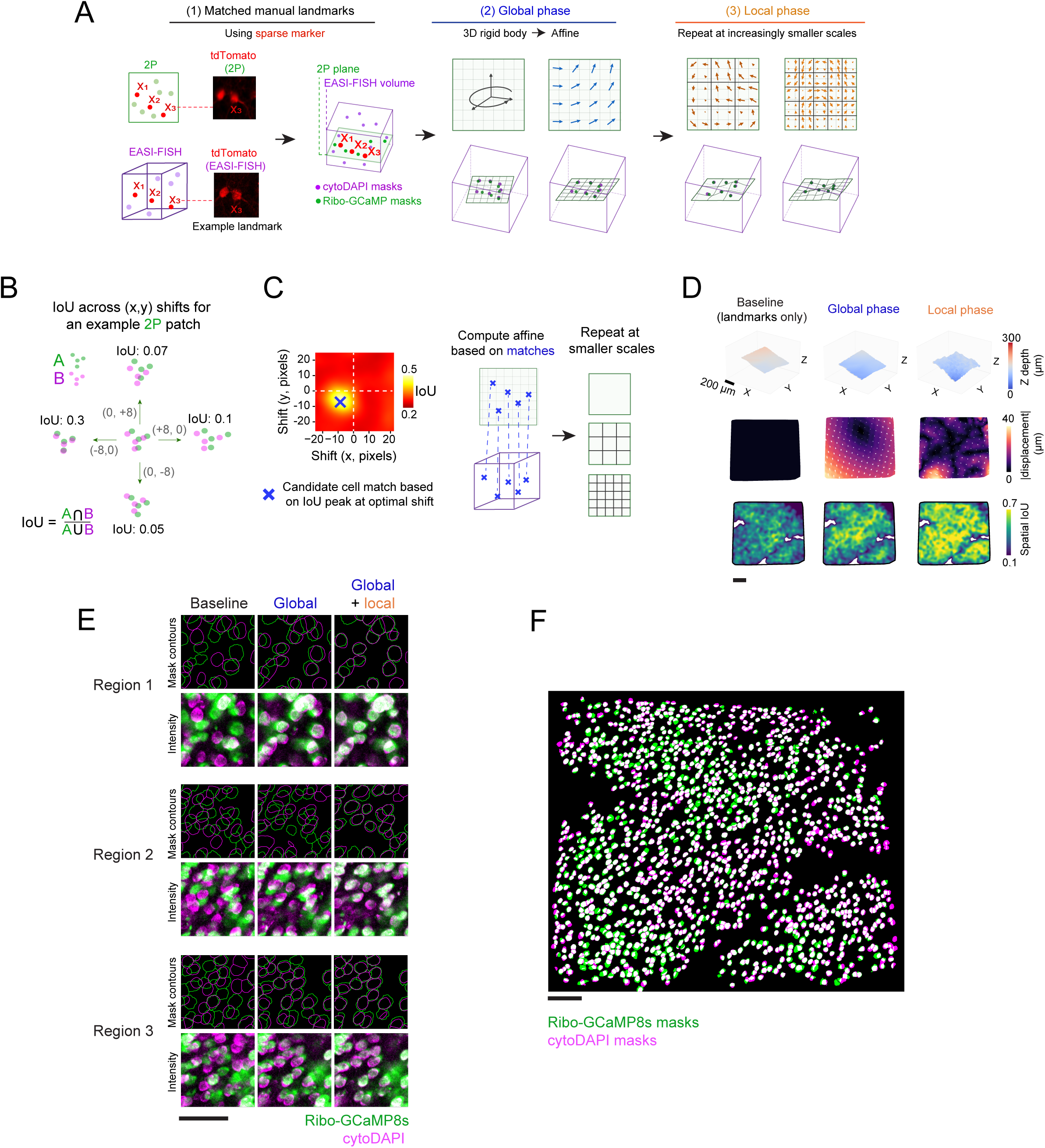
Alignment of 2P images and EASI-FISH volumes. **(A)** (1) Corresponding landmarks are placed in the 2P image and *ex vivo* EASI-FISH volume using a sparse anatomical marker (tdTomato). Landmarks are then fit with a thin-plate spline transformation to map the 2P plane onto a curved surface inside the EASI-FISH volume. (2) Global phase: a 3D rigid body search maximizes mask overlap (intersection over union, IoU). Cells whose masks overlap best under a local shift are taken as candidate matches and used to fit an affine transformation. (3) Local phase: to refine alignment, the above process is repeated for progressively smaller subregions, with iteratively updated candidate matches for local transformations. Green, 2P imaging plane; purple, EASI-FISH confocal volume. **(B)** IoU-based feature matching strategy for registration. Four example shifts for one 2P patch (green discs) mapped onto candidate EASI-FISH masks (purple discs) with the resulting IoU values. **(C)** Left: full map of IoU values across all shifts for one 2P patch. The peak identifies the best-aligned EASI-FISH cell. Right: matched cells are used to fit an affine transformation. The same procedure is repeated within progressively smaller subregions during local refinement. **(D)** Example 2P field of view, shown across baseline, global, and local alignment phases. Baseline indicates initial alignment using manual landmarks. Top row: 3D rendering of a single 2P plane within the EASI-FISH volume. Middle row: displacement field for each registration phase. Bottom row: spatially smoothed IoU between 2P and EASI-FISH masks. **(E)** 2P (green) and EASI-FISH (purple) mask contours and image intensity for three zoomed-in regions shown across alignment stages. **(F)** Full-field overlay of matched 2P and EASI-FISH masks for one plane. Scale bars, 100 µm unless otherwise noted.

### Cell matching across 2P and EASI-FISH

To identify matched cell masks between 2P images and EASI-FISH volumes, we first scored each 2P cell mask by its peak overlap (IoU value) with the best candidate EASI-FISH mask (**Figure 3A**). Deciding matches using the IoU metric alone requires users to select a threshold, and cells below the threshold could reflect either a genuine mismatch or a false exclusion due to residual local misalignment. To assign matches without a user-defined threshold, we used Soma-print^16^, which matches cells based on the geometric arrangement of their neighbors and calibrates each match against the distribution of second-best scores, resulting in a probability of correct registration for each cell (**Supplemental Figure 3A-B**). Soma-print and IoU-based matching (IoU > 0.3, see Methods) chose the same partner in ∼94% of cases when both methods matched the same cell. However, Soma-print matched ∼9% more cells than IoU-based matching, including a subset that fell below the IoU threshold (**Figure 3A; Supplemental Figure 3C-D**). We observed improved overlap across successive alignment stages when using either the Soma-print or IoU metrics, indicating that the alignment steps were important for increasing the number of matched cells and confidence in each match (**Figure 3B-C**). Relative to the landmark-only baseline alignment, the final alignment increased mask overlap, the fraction of cells matched by Soma-print, and the margin between the best and second-best Soma-print score in every sample (**Figure 3D-F**). We therefore used Soma-print (with minor adjustments; see Methods) to determine cell matches, and report both IoU and Soma-print metrics for every cell. In total, EASI-PASS matched 83% of 27,152 cells in the *in vivo* preparation (n = 5 mice) and 72% of 20,397 cells in the brain slice preparation (7 tissue slabs from 7 mice, **Figure 3G**).

**Figure 3.**
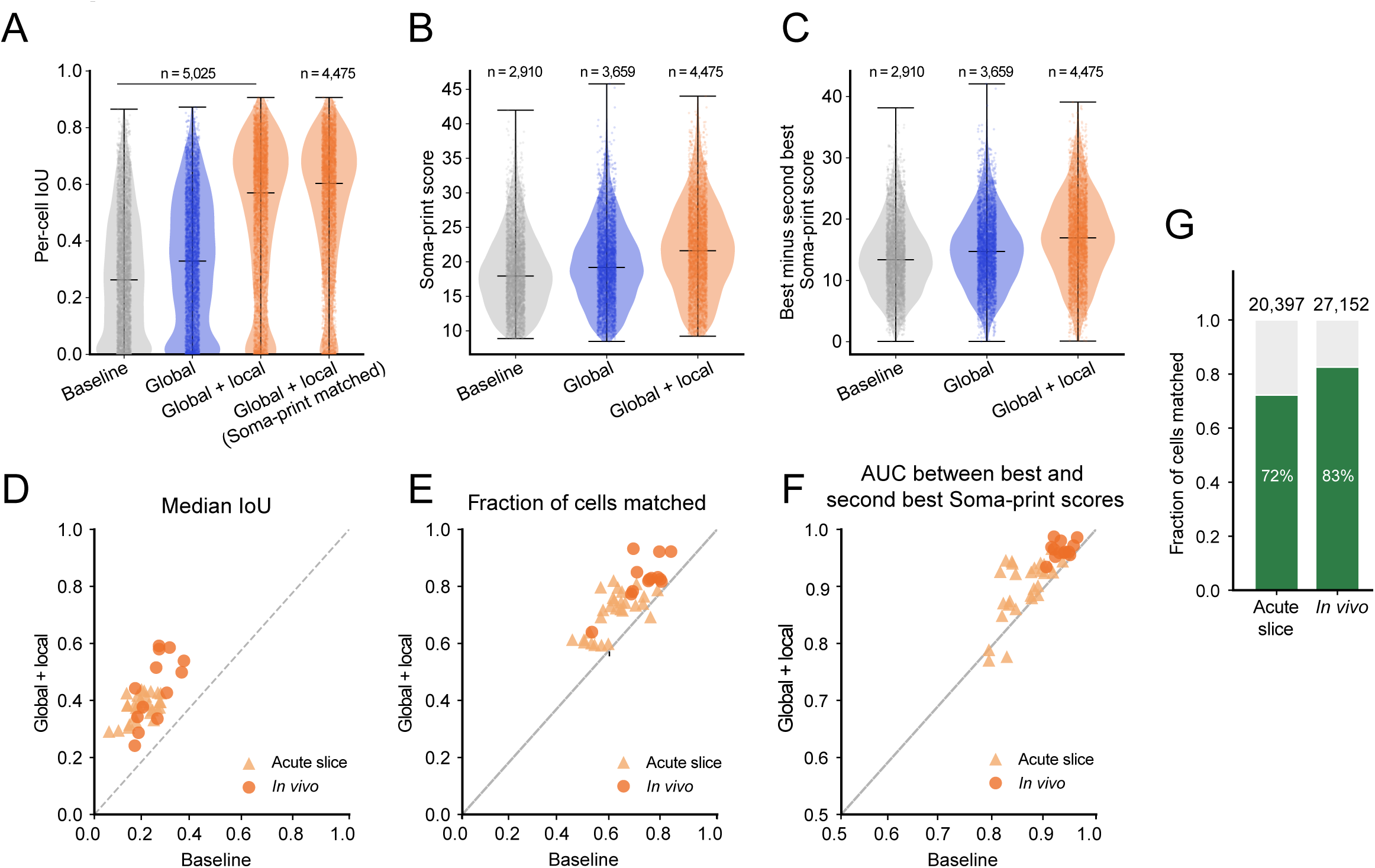
Cell matching across 2P and EASI-FISH. **(A)** Per-cell IoU across registration stages for one example mouse. The fourth column shows the same cells at the final stage, restricted to those matched by Soma-print. **(B)** Soma-print score of the best match for each confidently matched cell, across registration stages, for the mouse in (A). **(C)** Difference between the best and second-best Soma-print score, for the same cells as in (B). **(D)** Median IoU at baseline against median IoU after global and local alignment. Each marker is one 2P plane. Triangles, acute slice imaging. Circles, in vivo imaging. Two-tailed paired t test, baseline against global and local: acute slice, p = 1×10⁻⁵. *In vivo*, p = 0.002. **(E)** Fraction of cells matched by Soma-print, plotted as in (D). Acute slice, p = 0.001. *In vivo*, p = 0.002. **(F)** Discriminability between the best and second-best Soma-print scores, measured as the area under the receiver operating characteristic curve (AUC). Acute slice, p = 0.03. *In vivo*, p = 0.008. **(G)** Fraction of functionally imaged cells with a match, pooled across experiments. Acute slice imaging, 14,728 of 20,397 2P cells (72.2%). *In vivo* cortex imaging, 22,421 of 27,152 2P cells (82.6%).

### Spinoparabrachial input recruits a molecularly defined population of parabrachial neurons

We next turned to using EASI-PASS to uncover new biology in each of our preparations, starting with our acute slice imaging preparation of the PBN^31^. The PBN is a sensory hub for spinal, vagal, and humoral inputs^32–41^. At a coarse level, axonal inputs from these pathways show distinct spatial innervation profiles across PBN^35,36,42–45^ (see also **Supplemental Figure 4A**). Molecularly defined cell types in PBN are spatially clustered in various subregions, but with some degree of spatial overlap^46–48^. Together, these prior studies suggest a potential mapping of specific interoceptive pathways onto specific sets of cell types in PBN.

To directly examine this mapping, we focused on defining the PBN cell types activated by stimulation of spinoparabrachial axons from the lumbar dorsal horn, a key somatosensory pathway that relays multimodal signals, including noxious pain, itch, and temperature, to the rest of the brain^39–42,49–53^. Manipulations of this pathway are both necessary and sufficient for many pain-related behaviors^39–41,49,51,53,54^, but whether spinorecipient PBN neurons share a specific molecular signature is unknown. Identifying such a marker would be a useful first step toward targeting the pain-related population while sparing neighboring neurons that control other physiological functions, such as breathing^55–58^ and malaise^32,59–61^. We therefore expressed Ribo-GCaMP8s in PBN neurons and a red-shifted channelrhodopsin, ChrimsonR-tdTomato, in lumbar dorsal horn axons innervating the PBN (**Figure 4A-B**). Several weeks later, we cut acute brain slices and recorded PBN activity while optogenetically stimulating the ChrimsonR-expressing spinoparabrachial axons. Axons tagged with tdTomato were preserved after EASI-FISH and served as sparse markers for alignment. A low-magnification objective and a custom imaging chamber let us capture most of PBN in one field of view (1.2 mm^2^), so that we recorded from thousands of neurons (average 2,914 per slice) across several subregions at once (**Supplemental Figure 4B**). We observed spontaneous activity in most PBN neurons (**Figure 4C**). Consistent with the axonal innervation pattern, optogenetic stimulation reliably excited a subset of neurons almost entirely confined to the lateral PBN (LPBN, **Figure 4C-D**).

**Figure 4.**
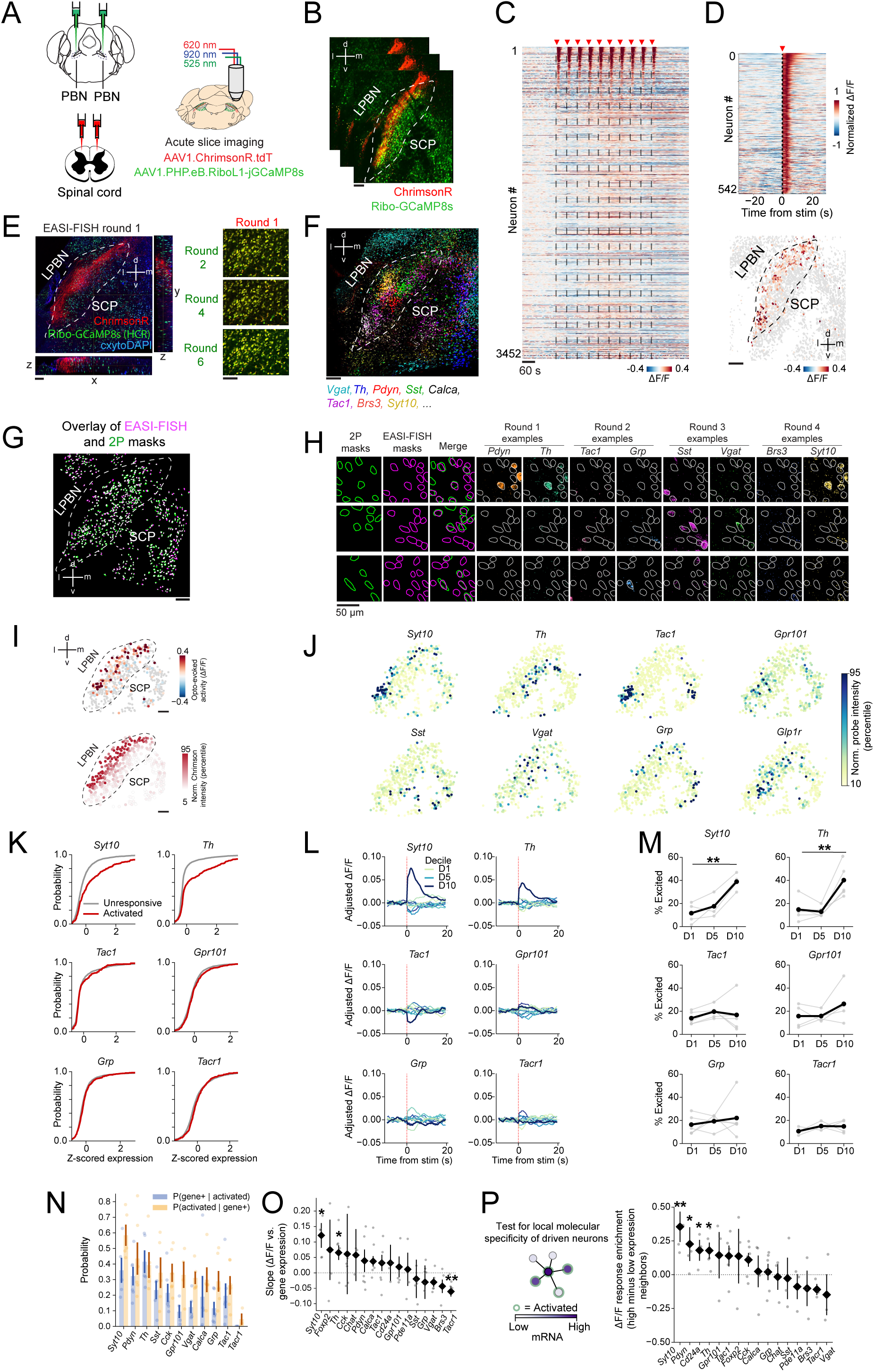
Spinoparabrachial input recruits a molecularly defined population of parabrachial neurons. **(A)** Experimental approach for acute slice two-photon calcium imaging of the parabrachial nucleus (PBN) during optogenetic stimulation of lumbar spinal cord axonal inputs expressing ChrimsonR-tdTomato. Soma-targeted Ribo-GCaMP8s reduced neuropil contamination. **(B)** Example imaging field of view. **(C)** Heatmap of neuronal activity for all neurons from the slice in (B) during 10 trials of optogenetic stimulation (red triangles; 3.6 s trains of 32 pulses at 8.89 Hz, 7 ms pulse duration, 8 mW, 40 s inter-trial interval). ΔF/F: fractional change in fluorescence. **(D)** Top: neurons significantly activated by optogenetic stimulation, sorted by response magnitude. Bottom: masks colored by response magnitude (average from 0 to 5 s after stimulus onset). **(E)** Left: confocal image of the first round of EASI-FISH for the tissue in (B). Rectangles at bottom and right indicate cross-sections through the depth of the EASI-FISH volume (XZ and YZ). Right: example zoom-in overlay of cytoDAPI from the first round of EASI-FISH (red) with the second, fourth, and sixth rounds (green). **(F)** Maximum-intensity projection of 8 example mRNAs from 6 registered rounds of EASI-FISH for the tissue in (B). **(G)** Matched 2P (green) and EASI-FISH (purple) masks for an example 2P plane. **(H)** Example insets showing 2P and EASI-FISH masks and mRNA labeling from four EASI-FISH rounds. **(I)** Matched mask centroids colored by normalized opto-evoked activity (top) or average ChrimsonR intensity in a 25 µm radius (bottom), each plotted as a percentile across cells. **(J)** Matched mask centroids colored by mRNA expression for a subset of genes. **(K)** Cumulative distributions of within-mouse z-scored gene expression for significantly activated (red) vs. unresponsive (gray) neurons (see STAR Methods), for six example genes. Mean expression, activated (n = 1,242 neurons) vs. unresponsive (n = 7,782 neurons). Per-mouse mean expression, significantly activated vs. unresponsive: two-tailed paired t test with Benjamini-Hochberg false discovery rate across genes (n = 6 mice); *Syt10,* p = 0.006, *Th,* p = 0.007. **(L)** Stimulus-locked ΔF/F by expression decile, plotted as each decile’s mean minus the mean across all binned cells, for one example slice. **(M)** Percentage of activated neurons per gene-expression decile (D1, D5, D10 shown; gray lines, individual mice; black line, mean across mice). Per-mouse % significantly activated, top vs. bottom decile (D10 vs. D1): two-tailed paired t test with Benjamini-Hochberg false discovery rate (n = 6 mice); *Syt10,* p = 0.007, *Th,* p = 0.012. **(N)** Blue: fraction of significantly activated neurons expressing each gene. Yellow: fraction of neurons expressing each gene that are significantly activated. Bars, mean across mice; error bars, SEM across mice; dots, individual mice. Only genes labeled in 4 or more mice are shown. **(O)** Regression slope of evoked ΔF/F on gene expression. Linear mixed-effects model over all matched cells (n = 9,024 neurons from 6 mice), with soma position and local ChrimsonR expression as covariates, and mouse as a random effect. Diamonds, fixed-effect estimate; error bars, one standard error of that estimate. Gray dots, per-mouse slope. Stars, significance after Benjamini-Hochberg correction across genes: *q < 0.05, **q < 0.01, ***q < 0.001. **(P)** Per-gene difference in evoked ΔF/F for each high-expressing neuron compared to its five nearest low-expressing neighbors (median neighbor distance: 45.3 µm). Diamonds, mean across mice; error bars, SEM across mice; gray dots, individual mice. Stars, significance from a linear mixed-effects model with mouse as a random effect, after Benjamini-Hochberg correction across genes: *q < 0.05, **q < 0.01, ***q < 0.001. Data are mean ± SEM across mice unless otherwise noted. Scale bars, 100 µm unless otherwise noted.

To map the molecular identity of spinorecipient PBN neurons, we performed iterative *in situ* hybridization labeling of up to 20 marker genes across seven rounds of EASI-FISH (3 *in situ* probes per round; 6 slices from 6 mice; **Figure 4E-H**). We chose marker genes with known enrichment in particular PBN subregions that together would allow us to tile the entire LPBN^46,47^. Some markers (e.g., *Pdyn*, *Calca, Brs3*) label a single PBN subtype, whereas others (e.g., *Tac1*, *Syt10, Tacr1*) span several subtypes. In some cases, slightly differing sets of genes were labeled, so we only analyzed genes examined in four or more mice. Because the gene panel targeted PBN, we excluded neurons from surrounding brain regions and restricted the analysis to PBN neurons (9,024 matched neurons from 6 slices). Of these neurons, 1,242 (13.7%) were activated by optogenetic stimulation of spinoparabrachial axons.

We next plotted the gene expression signature of optogenetically activated neurons and unresponsive neurons (**Figure 4I-K**). Expression levels of *Syt10*, *Th, and Pdyn* were higher in activated neurons than in unresponsive neurons, whereas expression of most other genes showed largely overlapping distributions between the two groups. To investigate the relationship of gene expression to responsiveness without thresholding, we sorted neurons by intensity of expression of each gene and averaged the stimulus-evoked response within each decile. We found that neurons in the highest decile of expression of *Syt10*, *Pdyn* or *Th* showed larger evoked responses than other neurons (**Figure 4L**). Accordingly, the fraction of activated neurons increased across deciles of gene expression (**Figure 4M**). In parallel, we classified neurons into binary categories using thresholds for significant gene expression and significant activation (see Methods). This allowed us to estimate the fraction of activated neurons that significantly express each gene (P(gene+|activated)) and the fraction of gene-expressing neurons that are activated (P(activated|gene+), **Figure 4N**). *Syt10* showed the highest probability in both cases, suggesting that it labels spinorecipient PBN neurons (likely composed of multiple functional subtypes^40,52,54,62^) with relatively high specificity.

Activated neurons could be enriched for a certain gene simply because a higher density of cells expressing that gene overlaps with the axon terminal field. Thus, while proximity was a reasonable predictor for activation, the cellular-level alignment made possible by EASI-PASS allowed us to ask whether, at a local scale (<50 µm), spinal input preferentially recruited neurons expressing a given marker. To assess local molecular specificity, we took two approaches: First, for each gene, we used multiple linear regression to predict a neuron’s evoked response (ΔF/F) as a function of (i) gene expression, (ii) the density of nearby ChrimsonR-expressing axons, and (iii) location on a smooth surface (quadratic fit) to capture any coarse gradients in evoked responses. Among all genes, *Syt10* remained the strongest predictor of evoked response magnitude, even when accounting for a neuron’s proximity to spinal axons and to other activated neurons (**Figure 4O, Supplemental Figure 4C,** and see Methods). Second, we compared each neuron with high expression of a gene (high-expressing neurons) to its nearest low-expressing neighbors within a 50 µm radius. *Syt10*+ neurons showed larger stimulus-evoked responses and were more likely to be activated than low-expression neighbors (**Figure 4P**). Together, these analyses suggest that spinoparabrachial inputs show fine-scale molecular specificity in their targeting of PBN neurons.

### Distinct arousal tuning and spatial correlations across cortical interneuron classes

We next applied EASI-PASS to investigate large-scale population activity of inhibitory interneurons in the mouse visual cortex. Cortical interneurons are powerful regulators of information processing^63–65^. Classes of interneurons exhibit somewhat distinct anatomical, electrophysiological, and molecular characteristics^66^. Recent work has sought to determine how the molecular heterogeneity of interneuron classes relates to their diverse functional properties. For example, by mapping functional imaging to spatial transcriptomics and immunolabeling^14,17,20,67,68^, prior studies have uncovered compelling differences in functional tuning across interneuron classes^69,70^. However, it has been challenging to address questions about population activity structure within each class due to low numbers of imaged cells of a given class per experiment, and/or comparisons of classes across mice rather than within the same experiment and field of view.

Due to its compatibility with large-scale volumetric calcium imaging and high-throughput cell matching, EASI-PASS is well-positioned to address questions about the behavioral modulation and functional connectivity of genetically defined classes. We therefore expressed Ribo-GCaMP8s pan-neuronally in the visual cortex of mice expressing tdTomato in *Sst*+ neurons (Sst-Cre;Ai14 mice; **Figure 5A**) and performed 2P calcium imaging of thousands of neurons during spontaneous behavior on a running wheel (3 planes spaced 40 µm apart in layer 2/3, each imaged at 5.2 Hz). The tdTomato signals served as an anchor for the initial coarse alignment using EASI-PASS (see also **Figure 2**), as well as to validate the specificity and selectivity of alignment in the case of *Sst+* neurons.

**Figure 5.**
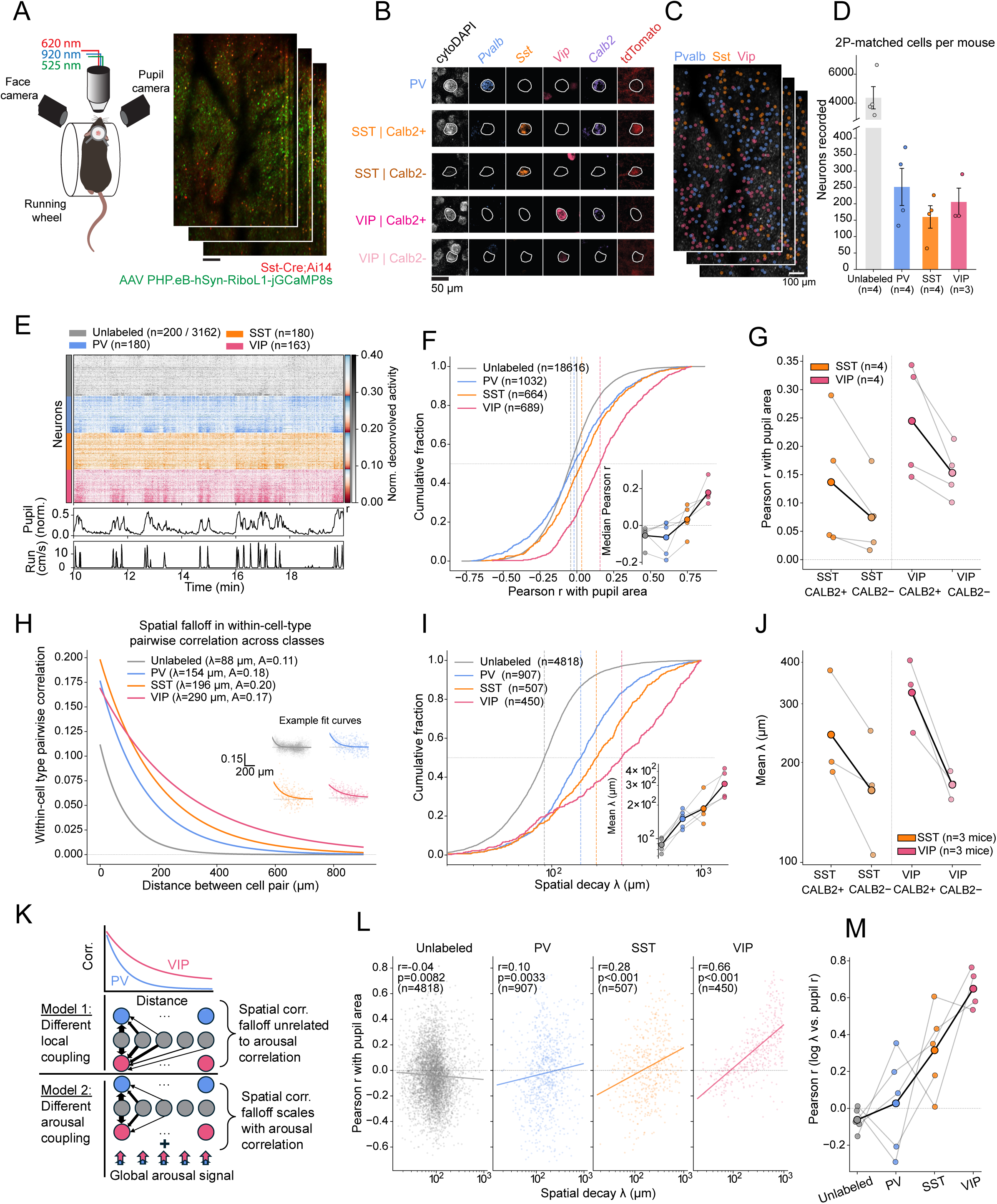
Molecularly defined cortical interneurons display graded differences in arousal tuning and spatial correlation structure. **(A)** Left: schematic of *in vivo* imaging preparation. Right: example field of view. **(B)** Zoom-in of EASI-FISH mRNA labeling for one example mouse, showing 4 genes as well as cytoDAPI, and tdTomato. **(C)** Same example field of view as in **(A)**, with cardinal interneurons labeled using the EASI-PASS pipeline. PV, blue; SST, orange; VIP, pink. **(D)** Numbers of matched neurons per cardinal class in mice for which all three imaged planes were contained in the EASI-FISH volume (4 of the 5 mice in this study). Unlabeled neurons are presumed to be excitatory. **(E)** Raster plot of deconvolved activity during spontaneous behavior in an example imaging session. Cells are sorted within class by Pearson correlation with pupil area (colorbar at right; red: positively correlated; blue: negatively correlated). Lower panels: traces of normalized pupil area and locomotion speed. **(F)** Cumulative distribution of Pearson correlation with pupil area, split by labeled cell class. Kruskal-Wallis within-mouse permutation test: H = 190.79, p < 0.001, n = 2,385 cells. Inset, Friedman test: χ² = 11.10, p = 0.011, n = 4 mice. **(G)** Per-mouse mean correlation with pupil area for subtypes of VIP or SST neurons, split by the presence or absence of *Calb2* expression. Within-mouse permutation test on the difference between *Calb2*+ and *Calb2*− subsets: SST, Δmean Pearson r = 0.062, p = 0.162, n = 600 neurons from 4 mice; VIP, Δmean Pearson r = 0.092, p < 0.001, n = 680 neurons from 4 mice. Gray lines, individual mice; black line, mean across mice. **(H)** Exponential decay curves derived from median fit parameters for within-class pairwise correlation in spontaneous activity as a function of distance between cells. Inset: example fits over data from four representative cells, one from each class. **(I)** Cumulative distribution of the exponential decay length constant (λ), split by cardinal class. λ differed across classes: Kruskal-Wallis within-mouse permutation test, H = 107.10, p < 0.001, n = 1,864 neurons. Inset, per-mouse median λ; Friedman test, χ² = 12.00, p = 0.007, n = 4 mice. **(J)** Per-mouse λ for VIP and SST neurons, split by *Calb2* expression. Within-mouse permutation test on the difference between *Calb2*+ and *Calb2*− subsets, computed on log λ: SST, Δmean log λ = 0.386, p < 0.001, n = 456 neurons from 3 mice; VIP, Δmean log λ = 0.638, p < 0.001, n = 360 neurons from 3 mice. Only mice with at least 5 successfully fit neurons per subtype are included. Gray lines, individual mice; black line, mean across mice. **(K)** Schematic of two toy models that could underlie observed differences across cell class in the falloff of within-class pairwise spontaneous correlations over distance. Model 1: VIP vs. PV cells have different falloff in coupling to local excitatory cells. Model 2: VIP cells have similar coupling to local excitatory cells but greater positive coupling to a global arousal signal, which effectively amplifies the correlation between spatially distant pairs of VIP cells. The two models make different predictions for the relationship between arousal coupling and λ within a class. **(L)** ë vs. pupil modulation (Pearson correlation with pupil area), plotted separately for cells in each cardinal class. **(M)** Within-class Pearson correlation between log λ and correlation with pupil area, for each cardinal class. Gray lines, individual mice; black line, mean across mice. Data are mean ± SEM across mice unless otherwise noted. Scale bars, 100 µm unless otherwise noted.

We then identified the imaged neurons with EASI-PASS and labeled major classes of interneurons with *Pvalb*, *Sst*, and *Vip* (**Figure 5B-C**). Because we imaged from multiple planes, each spanning ∼1.2 mm^2^, a single *in vivo* recording and corresponding EASI-FISH thick tissue yielded ∼150-250 simultaneously recorded interneurons of each class (PV, SST, VIP), together with ∼4,000 putative excitatory neurons per experiment (**Figure 5D**). As an independent validation of alignment quality, we quantified the overlap between tdTomato and our cardinal interneuron markers. We observed tdTomato expression in over 75% of EASI-FISH-labeled *Sst+* neurons, compared to 6.8% of *Pvalb+* and <1% of *Vip+* neurons (**Supplemental Figure 5A-B**), consistent with previous immunohistochemical characterization of this line^71^. As a further validation of our *in situ* signal, we compared the distribution of neuron depth within layer 2/3 across the cardinal classes and found the expected enrichment of VIP vs. SST neurons in superficial vs. deep layer 2/3, respectively^65,66^ (**Supplemental Figure 5C**). In the second round of EASI-FISH that we performed for this proof-of-principle experiment, we also assessed expression of *Calb2* to resolve subtypes within the *Sst* and *Vip* classes^66,72,73^. In future experiments, additional rounds of EASI-FISH will allow quantification of expression of several dozen other genes per neuron, as in **Figure 4**.

Prior work characterized how arousal states differentially modulate the activity of transcriptionally defined interneuron classes. Here, we first replicated these findings using large samples of neurons from each class in the same experiment, by quantifying how the activity of interneurons in each class covaried with pupil area, a sensitive correlate of arousal^74,75^ (**Figure 5E**). Consistent with prior studies, interneurons within each class showed diverse correlations with arousal, with a median correlation that differed across classes. Roughly equal subsets of *Pvalb*+ neurons were positively or negatively correlated with pupil area, while *Vip*+ neurons were predominantly positively correlated, and *Sst+* neurons showed an intermediate distribution^17,20^ (**Figure 5F**). The *Calb2*-defined subtypes of the SST and VIP classes showed distinct arousal profiles as well. Within each class, neurons co-expressing *Calb2* showed a higher positive correlation with pupil area, indicating higher activity in aroused states (**Figure 5G**).

We next asked how transcriptionally defined cell classes may differ at the level of within-class population activity. Taking advantage of our large imaging field of view, we investigated how different interneuron classes coordinate their spontaneous activity across distances exceeding the dendritic and axonal fields of most interneurons. Pairwise correlations between neurons of the same class located at different distances apart revealed that activity correlation decreases with distance (insets in **Figure 5H**). These correlation falloffs were well-captured using exponential fits. Pairs of putative excitatory neurons and pairs of PV neurons showed a sharp falloff with distance (**Figure 5H-I**). In contrast, VIP neurons remained correlated with other VIP neurons over long distances, and SST neurons exhibited an intermediate range of correlation falloffs with distance, mirroring the differences we observed in arousal coupling. Similarly, for pairs of VIP or SST neurons that expressed *Calb2*, pairwise correlations exhibited significantly larger spatial decay constants (**Figure 5J**), paralleling their increased levels of activity in aroused states. In contrast with falloff in correlations with distance, the different classes showed no significant differences in the intercept of the exponential fit (**Supplemental Figure 5D**), indicating relatively similar within-class correlations between nearest neighbors.

The above findings led us to consider two models of connectivity that could produce differences in spatial correlation structure across classes (**Figure 5K**). In both models, we assume that correlations between interneurons are partially driven by common inputs, such as shared local and/or long-range excitatory inputs (though spatially restricted cholinergic inputs^76^, gap junction coupling of nearby interneurons^77,78^, and other mechanisms are also likely to contribute). The first model posits that across-class differences in spatial falloff are driven by differences in the lateral extent of shared input to a pair of interneurons from the same excitatory neuron. For example, distant pairs of VIP neurons could be more correlated than distant pairs of PV neurons due to greater shared input from the same excitatory neurons (**Figure 5K, Model 1**). The second model posits that differences in falloff arise from differences in the contribution of shared arousal coupling to the spontaneous activity correlation between a neuron and its within-class neighbors independent of distance (**Model 2**). In this model, local input connectivity would be the same between VIP and PV neurons, but VIP neurons would exhibit greater correlation with distant VIP neurons due to the stronger and more consistently positive arousal modulation across VIP neurons (**Figure 5F**).

To distinguish which model better described our data, we assessed whether differences in a neuron’s long-range correlations (estimated using the spatial decay parameter, λ) are correlated with its arousal tuning. As predicted by the second model, VIP neurons with more gradual spatial falloff also exhibited stronger positive arousal modulation. However, this relationship was weaker in PV and SST neurons, and reversed sign in unlabeled neurons, despite a similarly broad range of arousal modulation and spatial decays for interneurons of each class (**Figure 5L-M**). This suggests that other factors (e.g., distinct integration of local excitatory input, as in **Model 1**) may have a stronger impact than arousal modulation on the falloff in pairwise correlations for these other interneuron classes. These findings were not simply due to variations in the mean falloff or arousal modulation of different subtypes within a given class, as similar trends were observed for both *Calb2*+ and *Calb2*-subtypes of SST and VIP neurons (**Supplemental Figure 5E**). We also ruled out that these findings might be due to differences in the quality of exponential fitting across classes (**Supplemental Figure 5F-H**). Taken together, these findings highlight how EASI-PASS provides a sensitive and efficient means to examine new biological questions requiring comparisons of the functional properties of multiple classes and subtypes of neurons in the same *in vivo* experiment.

## Discussion

Here, we describe a new pipeline, EASI-PASS, for cross-modal alignment of functional recordings and spatial gene expression in the same neurons. EASI-PASS builds on EASI-FISH, enabling thick-tissue mRNA profiling that retains sparse fluorescent markers and can robustly define the molecular identity of thousands of neurons in a single tissue slab. The pipeline does not require specialized hardware, finds matches for over three quarters of densely packed neurons per sample, and includes modules for segmentation, alignment across rounds of FISH, and quantification of mRNA expression. EASI-PASS runs largely automatically end-to-end, carrying raw image stacks through to a final table of matched neurons and their gene expression. EASI-PASS guides the user through the steps that require manual input or visual confirmation, such as landmark placement and confirmation of registration quality. In acute brain slice experiments, we established a calcium imaging approach for identification of neurons driven by long-range spinal inputs and identified a gene that defines spinorecipient PBN neurons with reasonable selectivity. In the visual cortex, we found that different molecularly defined interneuron classes and subtypes exhibit different degrees of falloff in pairwise correlations across space, a result that was partially but not fully explained by differences in arousal modulation.

Using our acute brain slice preparation, we recorded PBN neuron responses to optogenetic stimulation of spinoparabrachial inputs and observed a spatially restricted pattern of excitation. The large field of view (1.2 mm²) allowed us to ask how spinal inputs recruited neurons across the entire PBN and to profile ∼20 marker genes spanning all PBN subnuclei. In the future, this approach could be applied to genetically defined subsets of inputs from the spinal cord and other pathways that relay more specific body signals such as those related to pain, itch, temperature, or satiety. We found that spinorecipient neurons were enriched for *Syt10*, a calcium sensor involved in exocytosis^79^. *Syt10* labels the dorsolateral PBN, in agreement with previous work indicating the presence of spinorecipient PBN neurons in this subregion^54^. *Syt10* (and other markers biased to the dorsolateral PBN) predicted spinorecipient neurons better than genes enriched in other regions of lateral PBN such as *Tacr1* or *Calca,* both of which were suggested as recipients of spinoparabrachial input in previous studies^40,42^. Our alignment also allowed us to compare each gene-expressing neuron with its nearest neighbors, a direct test of local molecular specificity. *Syt10*+ neurons tended to be more strongly driven than immediate neighbors with low *Syt10* expression, suggesting that the bias towards *Syt10*+ neurons is also present at a very fine scale. This finding highlights the importance of precise matching of functional properties with molecular expression at the cellular level, even for regions such as the PBN where many molecular cell types show local spatial clustering. While the combination of optogenetic stimulation of long-range inputs with dense calcium imaging allows high-throughput assessment of sensitivity to spinoparabrachial inputs (**Figure 4**), we cannot confirm that the activation is due solely to monosynaptic spinoparabrachial connections. However, future experiments could assess such monosynaptic connectivity by combining EASI-FISH with anterograde transsynaptic tracing^80–82^.

By applying EASI-PASS to the visual cortex of awake, behaving mice, we could routinely perform simultaneous recordings in large populations of neurons from each of the three cardinal classes of inhibitory interneurons (>150 neurons per class, on average). This high yield will be critical in studies attempting to make quantitative statements about increasingly rare subtypes of each interneuron class, such as those defined here by the presence or absence of *Calb2* expression. The large spatial scale of these recordings also enabled population-level analysis of simultaneously recorded, molecularly defined interneurons, revealing a new, simplifying description of the organization of correlated spontaneous activity over space: interneuron classes and subtypes with more consistently positive arousal modulation coordinate their within-class spontaneous activity over larger spatial scales. This coupling between higher arousal modulation and weaker falloff in correlated spontaneous activity was stronger for VIP neurons than for other classes. This suggests that factors other than arousal (e.g., local pyramidal neuron coupling, visual tuning properties, gap junction coupling and/or sensitivity to locally released neuromodulators and peptides) may explain a greater degree of variability in the spatial falloff in coupling between a PV neuron or an SST neuron to other neurons of the same class.

Our results also emphasize the importance of subtype markers in describing cortical interneuron function *in vivo*. For both SST and VIP neurons, *Calb2* expression explained some of the within-class variance in the arousal modulation of each neuron and the spatial falloff of spontaneous correlations. Despite the fact that expression of *Calb2* predicts differences in morphological properties across SST subtypes (degree of axonal bias toward layer 1^66,72^) and across VIP subtypes (degree of axonal bias toward L5^73^), both SST/Calb2+ and SST/Calb2-subtypes are similarly enriched in expression of excitatory cholinergic receptors^17^, which may help explain their increased arousal modulation.

Several methods have recently emerged for cross-modal image alignment, yet their adoption is often limited by the need for specialized equipment and the low numbers of matched cells recovered. Some of this loss (and additional labor) comes from the challenge of aligning 2P images to many thin *ex vivo* sections. In contrast, EASI-PASS registers a functionally imaged plane into an intact 3D EASI-FISH volume. Our approach recovers thousands of matched neurons per experiment with high yield, requires no specialized equipment, is computationally inexpensive, and is openly available. Nonetheless, important limitations remain. The initial alignment requires the user to place landmarks by hand (in our case, guided by a sparse marker) which can be time-consuming if accurate records of tissue orientation are not maintained throughout post-hoc processing stages (with accurate records, this process usually takes less than 30 minutes). Registration quality also depends on uniform expansion and complete clearing during EASI-FISH^21,22,83^, so users should ensure these steps are reliable for their applications, which may require adapting the protocol to various brain tissues. Finally, we summarized each cell’s molecular signal as a mean fluorescence intensity rather than using a higher-magnification objective to allow counting of individual mRNA puncta. We made this choice to prioritize a combination of increased throughput and coverage together with reduced data acquisition time, data storage and overall cost. While our approach is adequate when using high-abundance genes to define cell types, puncta counting can be readily integrated where single-molecule resolution is needed (e.g., for lower-abundance genes such as those for G-protein coupled receptors). By linking activity to molecular identity at scale and with minimal specialized equipment and manual labor, EASI-PASS should help neuroscientists address a broad range of new questions, such as how the coordinated activity of many simultaneously recorded, molecularly defined cell types underlies circuit functions and behavior.

## Acknowledgments

We thank A. Haydaroğlu, R. Essner, G. Chahyadinata and members of the Andermann lab for useful feedback. We also thank B. Lowell, S. Nardone, and R. Grippo for transcriptomic sequencing expertise and wet lab assistance in early versions of the protocol. A. Chirila for technical assistance with spinal cord viral injections. J. Campos, M. Garrett, P. Groblewski for assistance on cortical window tissue preparation. G. Fishell for advice on cortical gene panel design. A. Jaggi for consultation on durotomy procedure. H. Li, A. Pinilla, and J. DeBolt for helping with animal care, behavioral experiments, and reagent acquisition. Boston Children’s Hospital Viral Core (NIH P30 EY012196) and HMS Research Instrumentation Core provided viral and laser printing services, respectively. Authors were supported by a Harvard Brain Initiative Postdoctoral Pioneer Fellowship, a Harvard Mind Brain and Behavior Postdoctoral Fellowship, and NIH K99 NS140561 (J.S.A.); an Albert Ryan Fellowship and an Ellen R. and Melvin J. Gordon Fellowship in Neurobiology (C.M.); an NIH NRSA F30 DA057823 and NIH T32 GM007753 (K.E.E.); an NIH DP1 AT010971, R01 MH123431, R01 EY032749, R01 EY013613, R21 EY035436, DP1 DK139958, Pew Innovation Grant, the Nancy Lurie Marks Foundation, The Harvard Brain Science Initiative Bipolar Disorder Seed Grant supported by Kent and Liz Dauten, the K. Lisa Yang Brain Body Center, and a gift from Paul Weisman (M.L.A.).

## Author contributions

Conceptualization, J.S.A., O.A. and M.L.A.; investigation, J.S.A., C.M., J.S., B.V., S.C., A.V., P.S., D.P., J.C., K.E.E., A.J.; formal analysis and visualization, J.S.A., J.S., O.A., C.M., and M.L.A.; writing – original draft, J.S.A., J.S., M.L.A.; writing – review & editing, J.S.A., J.S., Y.W., and M.L.A.

## EXPERIMENTAL MODEL AND SUBJECT DETAILS

### Mice

All animal care and experimental procedures were approved by the Beth Israel Deaconess Medical Center Institutional Animal Care and Use Committee. Mice were group housed before surgery and singly housed afterward, with ad libitum access to standard chow and water, under a 12 h–12 h dark–light cycle at 20 to 22 °C and 30 to 70% humidity. All experiments were performed during the light cycle.

Cortical imaging was performed in adult (older than postnatal day 56) *Sst*-Cre;Ai14 mice, which express Cre recombinase in somatostatin interneurons together with a Cre-dependent tdTomato reporter. PBN slice imaging with spinal afferent stimulation was performed in wild-type mice aged 9 to 15 weeks. Mice of both sexes were used for cortical imaging. Only male mice were used for the parabrachial slice experiments with spinal afferent stimulation, since females at postnatal day 21 are too small for spinal injection.

## METHOD DETAILS

### Stereotaxic injections

Surgery was performed under isoflurane anesthesia (1.5%) on a stereotaxic frame (Kopf Instruments, Model 940), with body temperature maintained on a heating pad. For cortical imaging, AAV1-PHP.eB-hSyn-RiboL1-jGCaMP8s virus was diluted to 1 × 10^12^ gc/ml and delivered through the craniotomy described below at three to six sites, evenly spaced across the exposed circular brain surface, with three depths per site (250, 350 and 500 µm below the pia) and 100 nL per depth at 20 nL/min. For acute slice imaging, AAV1-hSyn-ChrimsonR-tdTomato was injected bilaterally into 2-3 sites (50 nL per depth at 20 nL/min) in the lumbar spinal dorsal horn to label ascending spinoparabrachial axons. After waiting one week for recovery, AAV1-PHP.eB-hSyn-RiboL1-jGCaMP8s was injected bilaterally into the parabrachial nucleus (AP −5.2 mm, ML ±1.5 mm, DV −3.85 & 3.55 mm from Bregma, 100 nL per depth at 20 nL/min). Mice were allowed at least 5 weeks for expression before acute slice imaging experiments were performed.

### Cranial window surgery

Cranial windows were implanted following previously described procedures^1^. Briefly, in anesthetized mice, a 3 mm circular craniotomy was made with a dental drill over the higher visual areas of the left hemisphere (AP +1.55 mm from the caudal sinus, ML −4.3 mm from Bregma), and the skull surrounding the craniotomy was thinned. We found that removing the dura was necessary for full integration of the tissue into the EASI-FISH gel. This was done either by durotomy at the time of window implantation or, where no durotomy was performed, by trimming 20 to 30 µm from the pial surface of the tissue block after extraction (see Recovery of the imaged cortical tissue, below). Virus was injected across the exposed brain surface as described above. A double-layer window was then placed on the brain surface, consisting of a 3 mm circular coverglass bonded to a 5 mm coverglass. The smaller glass sat within the craniotomy against the cortex while the edges of the larger glass rested on the thinned skull, which holds the window flush and stabilizes the tissue. The window was fixed in place with C&B Metabond (Parkell), and a titanium headpost was affixed to the skull. Mice recovered for at least 4 weeks before imaging.

### Retinotopic mapping

Visual areas were mapped by widefield epifluorescence imaging of Ribo-GCaMP8s. Excitation was provided by a 470 nm LED, emission was collected through a 500 nm long-pass filter, and images were acquired with an EMCCD camera while low-contrast vertical and horizontal bandpass noise was presented at four retinotopic positions on a 60 Hz LCD monitor positioned to the right of the mouse. The resulting maps of local preference for position in visual space were used to identify V1, LM, AL, RL and LI and to place the 2P field of view. Area boundaries were drawn manually after alignment to a reference atlas, and each recorded cell was assigned to an area based on the center of mass of its cell mask.

### In vivo 2P imaging

Multi-plane imaging was performed on a resonant-scanning 2P microscope (Neurolabware) with a 16×, 0.8 NA water-immersion objective (Nikon), an InSight X3 laser (Spectra-Physics) and an electrically tunable lens (Optotune). Ribo-GCaMP8s was excited at 920 nm and tdTomato at 1100 nm. Layer 2/3 of lateral V1, AL and LM was recorded for 30 to 60 min per session in the absence of visual stimulation. The field of view was 0.8 × 1.2 mm (796 × 512 pixels). We imaged three planes spaced 40 µm apart at 5.21 frames per second per plane, spanning approximately 100 to 220 µm below the pial surface.

### Behavioral monitoring during imaging

At least 3 weeks after surgery, mice were habituated to head fixation over 4 to 7 days. On the first day, mice were head-fixed for 30 minutes and allowed to run on a wheel in darkness. For each subsequent day, the training session lasted 1.5 hours. A gray screen was presented on the LCD monitor from the second session onward. Habituation continued until a mouse remained calm for the full session, with no physical signs of stress.

During imaging, mice were head-fixed on the wheel, which was locked in a subset of sessions and free to rotate in the other sessions, allowing postural adjustment in both cases. The right eye and the left side of the face were each recorded with infrared cameras. Pupil and eyelid keypoints were tracked with DeepLabCut^2^, using a custom model trained on approximately 1,000 manually labeled frames with eight points around the exposed eye.

### Acute slice preparation and imaging

#### Slice preparation

Mice were deeply anesthetized with isoflurane and decapitated. Brains were extracted and immediately immersed in ice-cold, carbogen-saturated (95% O2, 5% CO2) choline-based cutting solution containing (in mM) 92 choline chloride, 10 HEPES, 2.5 KCl, 1.25 NaH2PO4, 30 NaHCO3, 25 glucose, 10 MgSO4, 0.5 CaCl2, 2 thiourea, 5 sodium ascorbate and 3 sodium pyruvate, at 310 to 320 mOsm/L and pH 7.4. Coronal sections (275 µm thick) containing the PBN were cut on a vibratome (Campden 7000smz-2) and recovered in oxygenated cutting solution at 35 °C for 15 min, then in oxygenated aCSF containing (in mM) 126 NaCl, 21.4 NaHCO3, 2.5 KCl, 1.2 NaH2PO4, 1.2 MgCl2, 2.4 CaCl2 and 10 glucose at 35 °C for 30 min. Slices were brought to room temperature for at least 45 min before recording. A single slice was transferred to a custom imaging chamber and continuously superfused with carbogen-saturated aCSF at 2 to 5 mL/min at room temperature.

#### Imaging

To capture the entire PBN and its surroundings, we used a 10×, 0.6 NA water-immersion objective (Olympus XLPLN10XSVMP, 1.2 mm^2^) so that all PBN subregions were recorded simultaneously. Planes were acquired with an electrically tunable lens (ETL; Optotune). Ribo-GCaMP8s was excited at 920 nm. We imaged four to five planes over a 796 × 900 pixel field of view at 3 to 5 frames per second per plane. Planes were spaced 30 to 40 µm apart, beginning 15 to 30 µm below the surface of the slice. ChrimsonR-expressing spinal inputs were activated with a 620 nm LED (Luxeon Star) positioned beneath the slice and driven by a programmable Arduino controller, to deliver wide-field illumination (8 mW). Each trial consisted of a single train of 32 pulses of 7 ms duration at 8.89 Hz. Stimulation was synchronized to frame acquisition, with the photomultiplier tubes (PMTs) gated off during each pulse. Ten trials were delivered per recording, with an inter-trial interval of 40 seconds.

#### Fixation

After imaging, slices were drop-fixed in 4% PFA for 20 min at room temperature. Immediately after, slices were washed in 1× PBS (3 × 5 min) and stored in 70% EtOH until processing for EASI-FISH.

### High-resolution imaging

Manual landmarking requires identifying the same cell in the 2P image and EASI-FISH confocal volume. Finding matching landmarks is challenging at low resolution. Thus, when functional imaging experiments sampled cell bodies at fewer than 7 pixels per cell diameter, these experiments were followed by a high-resolution image of the same field of view. Magnification was increased so that each cell spanned at least 21 pixels in diameter, and overlapping tiles (20% overlap) were acquired at 920 nm and 1100 nm until the full functional field of view was covered, following by automated digital stitching of these higher-resolution images. The stitched high-resolution images were used to facilitate manual landmarking.

### Recovery of the imaged tissue in awake mouse cortical imaging experiments

#### Extraction

Mice were transcardially perfused with 4% PFA within 24 h of the final imaging session. Brains were extracted from the ventral side, post-fixed in 4% PFA for 3 to 6 h at room temperature, transferred to 20 °C for an additional 12 hours, and switched to 1× PBS. The imaged region was isolated with three razor cuts, along the midline and parallel to the anterior and posterior edges of the cranial window, whose imprint on the tissue was visualized under oblique illumination and used as the physical reference for the cutting angle.

#### Embedding

The tissue block was inverted onto a glass slide with the pial surface flush against the glass and centered within a 3D-printed ring. Low-melting-point agarose (4%) was applied until a dome formed above the ring, and a second slide was laid over it to flatten the surface. Once set, the assembly was inverted and the upper slide removed, leaving the tissue embedded except at the cortical surface beneath the window, which was now parallel to the agarose surface and thus parallel to the vibratome blade (see below). A two-photon z-stack spanning 500 µm from the surface of the embedded block was acquired. This stack was later used to locate the imaged field during trimming, by matching the pattern of blood vessels and of bright fluorescence at the viral injection sites against the same features viewed under the fluorescence microscope (see Trimming and orientation for EASI-FISH, below).

#### Sectioning

The tissue block was then mounted on and sectioned on a vibratome (Leica VT1000 S). The surface was approached in 10 µm steps until first contact with the tissue was observed. From here, an additional 30 µm was cut to clear the dura and superficial scar tissue, and a final 200 µm section containing the imaged field was collected into 1× PBS for use the next day, or 70% EtOH for long-term storage.

### Trimming and orientation for EASI-FISH

Cortical sections were placed under a fluorescence microscope (Olympus SlideView VS200). The imaged region was located from blood vessel patterns and from bright fluorescence at the viral injection sites. Acute slices were viewed under brightfield illumination instead, and the PBN was located from anatomical landmarks, including the superior cerebellar peduncle and the cerebellum. Tissue was then trimmed manually under a dissection microscope with a fresh razor blade, using anatomical landmarks as boundaries. The final area was approximately 2 mm². A diagonal corner cut was made to preserve orientation through the remaining steps (**Supplemental Figure 1A**). Tissue was trimmed on a glass slide with the imaged surface facing upward, away from the glass. Tissue was placed in the same orientation for gelation. In this way, the tissue would sink to the bottom of the gel during formation, with the functionally imaged side facing the gel interior. This strategy was used to protect the imaged side from mechanical damage during handling. Samples were trimmed again after clearing and expansion (see below).

### EASI-FISH protocol modifications

For the majority of steps, we followed the published EASI-FISH protocol^3^. In some cases, we made modifications, which are described below.

#### mRNA anchoring

Trimmed samples were placed in MelphaX and incubated overnight (37 °C, 16 h) to anchor mRNA before gelation (Wang et al., 2021). MelphaX was prepared by reacting melphalan (2.5 mg/mL in anhydrous DMSO) with Acryloyl-X (10 mg/mL in anhydrous DMSO) at 4:1 (v/v) and applied 1:1 with MOPS buffer (20 mM). The combination is exothermic, so the solution was left to rest for 10 min at room temperature before use. Samples were then exchanged into 1× PBS by replacing half the volume five times, followed by two 10 min washes in 1× PBS. We found that these two modifications helped preserve tdTomato fluorescence.

#### Gelation, digestion and expansion

A glass slide was precoated with poly-L-lysine and allowed to air dry for 10 minutes. Samples were placed in a 9 mm diameter × 0.5 mm deep gasket (Invitrogen) on the coated slide. We modified the original protocol by first incubating the tissue in inactivated monomer solution (Stock-X, 30 min, 4 °C) before equilibrating in activated gel solution (40 µl, 3 × 10 min, 4 °C). We found that this step improved clearing and increased cytoDAPI brightness. The gel solution is composed of Stock-X activated with 4-hydroxy-TEMPO (0.5%), TEMED (10%) and APS (10%) at 94:2:2:2, made fresh and kept on ice. The gasket was sealed with a coverslip and the gel formed at 37 °C for 120 min. We ran this incubation in a humidified chamber (a glass Pyrex container lined with damp paper towels and pre-warmed to 37 °C), because the gel otherwise dries at the exposed gasket edge and tears when the coverslip and gasket are lifted off to recover the tissue-gel. Recovered gels were trimmed with a corner cut for orientation, digested overnight (37 °C) in Proteinase K buffer (750 µl) with Proteinase K (800 U/ml, 7.5 µl), then trimmed again and washed in PBS (4 × 15 min).

#### Photobleaching

Tissue samples collected from mice older than 8 weeks exhibit increased autofluorescence from lipofuscin spots. To mitigate this, samples underwent a 24-hour photobleaching treatment after gelation and clearing, as follows. After trimming and washing, expanded gels were placed in an uncovered well plate filled with 1× PBS. The plate was placed inside a photobleaching box (an uncovered plastic box lined on all surfaces with black light-blocking fabric) positioned 12 inches beneath a 200W LED light (SolarXtreme® 250, California Lightworks, 120 V), inside a 4 °C walk-in cold room. The box and light assembly were then fully covered with an additional sheet of light-blocking fabric. Once in the covered photobleaching setup, samples were bleached for 24 h, with PBS topped off every 3 hours to prevent drying.

#### DNAse I digestion

Following photobleaching, the LED was switched off and samples were digested in DNase I (QIAGEN, 2.7 Kunitz units/µL) diluted 1:10 into home-made DNase I buffer (10 mM Tris-HCl pH 8, 2.5 mM MgCl₂, 0.5 mM CaCl₂), 250 µL per well in a 48-well plate, for 2 hours at 37 °C with shaking. DNAse I digestion was performed twice (total 2 x 2 hours) before the first round of HCR^3^ (see HCR section below for details).

#### Sample preparation for confocal imaging

A reusable imaging chamber was assembled on a glass slide from two stacked plastic frames (SunJin Lab, 0.5 mm) bonded with optical-curing epoxy. The glass slides were coated with poly-L-lysine and air dried for at least 15 min. Poly-L-lysine holds the gel in place for each imaging session. Before each round of imaging, gels were stained with DAPI (0.5 µg/mL in PBS, 30 min) and washed in PBS (3 × 15 min) on a shaker. DAPI binds cytosolic RNA as well as nuclear DNA. Because DNase digestion removes nuclear DNA, DAPI staining after digestion labels cytosolic RNA, yielding a cytosolic stain (cytoDAPI) used for segmentation and registration^3^. The gel was gently placed on the slide with the tissue-side facing the objective (meaning the imaged side of the tissue is facing away from the slide, into the gel). The chamber was filled with PBS and sealed with a coverslip. No additional sealant is used; the seal is held through liquid adhesion between the coverslip and PBS. Care was taken to eliminate formation of bubbles in the imaging chamber, as these can lead to movement of the sample during long imaging sessions.

#### Confocal imaging

EASI-FISH volumes were acquired on a Leica SP5 confocal microscope with a 10×, 0.40 NA air objective (HC PL APO). Lateral sampling was 0.8 to 1.3 µm and axial steps were 5 to 7.5 µm (0.5 to 0.8 µm and 3 to 4.5 µm before expansion, given the EASI-FISH 1.7-fold expansion factor). Imaged volumes covered 1.5 to 3.0 mm² in area and 400 to 600 µm in thickness after expansion. Acquisition required 1.5 h for 5 channels and 4 tiles, with 10% overlap. In each round, cytoDAPI was imaged at 405 nm and mRNA probes at 488, 647 and 750 nm. The 546 nm channel was reserved for the sparse red label and left unprobed in every round. tdTomato fluorescence remained stable across 7 EASI-FISH rounds.

### Hybridization chain reaction and multi-round imaging

#### Stripping

Probes and hairpins from the previous round were removed by DNase I digestion. Gels were equilibrated in DNase I buffer (shaker, 30 min, 37 °C), then digested in DNase I (QIAGEN, 2.7 Kunitz units/µL) diluted 1:10 into home-made DNase I buffer (see Photobleaching section for recipe), 250 µL per well in a 48-well plate, for 120 min at 37 °C with shaking. Two consecutive digestions are performed before the first HCR round to initially produce cytoDAPI, but a single digestion is sufficient in later rounds for probe stripping. Gels were then washed in PBS (shaker, 4 × 15 min) to prevent residual DNase activity.

#### Hybridization

Gels were equilibrated in hybridization buffer (300 µL, 30 min, 37 °C) and hybridized overnight at 37 °C with the three probe sets for that round (HCR v3.0^4^, Molecular Instruments; 1 µM each,1 µL per 100 µL of tissue section hybridization buffer). Unbound probe is washed out in probe wash buffer (2 × 15 min, then 2 × 30 min) and PBS (5 × 15 min), both at 37 °C on a shaker.

#### Amplification

Gels were equilibrated in amplification buffer (200 µL per well, 30 min, room temperature). Hairpins carrying the three fluorophores were snap-cooled (95 °C, 90 s, then 30 min in the dark at room temperature), diluted 1:100 into amplification buffer, and placed on a shaker at room temperature for 3 hours. Unbound hairpins were washed out in 5× SSCT (5× SSC, 0.1% Tween; 2 × 20 min), then 0.5× SSCT (2 × 20 min) on a shaker at room temperature.

### Fluorescence time course extraction from 2P calcium imaging

All 2P data were motion-corrected in Suite2p^5^. Cell masks were obtained by segmenting the mean of the motion-corrected volume with Cellpose4^6^. Fluorescence was then measured frame by frame as the mean of all pixels within each mask. Because Ribo-GCaMP8s is confined to the soma, traces were analyzed without neuropil correction.

#### Slice recordings

Traces were converted to ΔF/F trial by trial as (F − F0)/F0, with F0 the mean fluorescence over the 10 s preceding each stimulus train.

#### Cortical recordings

F0 was taken as the 10th percentile of each cell’s trace over the full recording. ΔF/F was computed as above. Calcium events were deconvolved with OASIS^7^ using a decay time constant of 0.6 s and the per-session frame rate. The 99th-percentile-normalized deconvolved rate, smoothed with a Gaussian kernel (σ = 500 ms), was used for all cortical analyses.

### EASI-PASS pipeline

#### Overview

The EASI-PASS pipeline runs in five stages. Each EASI-FISH round is a confocal volume stack of the expanded tissue. (i) Cells are segmented (Cellpose4^6^) in each EASI-FISH round and in the 2P mean image of each functional plane. The first acquired EASI-FISH round serves as the reference volume onto which 2P and additional EASI-FISH rounds are registered. (ii) Each 2P plane is warped into the reference EASI-FISH volume. The warp is initialized from landmarks placed by hand in BigWarp, then improved by a 3D rigid body search, a global affine transform, and several additional affine transformations at progressively smaller spatial scales. Each registration stage maximizes the intersection over union (IoU) between 2P and EASI-FISH masks. We defined IoU as the number of pixels shared by the two masks (the intersection), divided by the total number of pixels covered by either mask (the union). When a high-resolution intermediate image is acquired, the functional image is first registered to that image, and the two transformations are then applied in sequence. (iii) 2P and EASI-FISH cells are matched using Soma-print^8^ or IoU; both metrics are reported for every cell. (iv) Additional EASI-FISH rounds are registered and matched to the reference volume. (v) Per-cell mRNA fluorescence and the match assignments are combined into the output table, with one row per cell of the reference round.

#### Inputs and workflow

EASI-PASS takes as inputs a two-dimensional mean image of the functional field of view and a three-dimensional volume for each FISH round, both as TIFF files. The pipeline is modular: a higher-magnification image of the functional field (see High-resolution imaging section) can be supplied as an alignment intermediate, and the pipeline can also run on multi-round FISH data alone, with no functional imaging. Each sample is configured by a single text file containing all input paths and pipeline parameters (see https://github.com/orena1/easi-pass for documentation). User input is required at three points: placement of the “cross-modal” landmarks in the 2P images and the corresponding locations in the initial EASI-FISH volume (in BigWarp^9^), curation of the 2P mask segmentation process, and verification of the round-to-round EASI-FISH alignment. Every other step is automated. Parameter values quoted below are the defaults we used and are intended as a starting point (see documentation for more details).

#### Cell segmentation

EASI-PASS segments cells with Cellpose4^6^. The model and evaluation parameters are specified per modality in the sample’s configuration file, and no downstream step is tied to a particular model or parameter set. The values below should be adjusted to fit each user’s needs. EASI-FISH volumes are segmented plane by plane in two dimensions and assembled into three-dimensional cells by linking masks between adjacent planes by overlap (do_3D=False, stitch_threshold = 0.3). Masks with outlier volumes (<0.1× or >10× the median mask volume) are then discarded. For the samples reported here, we segmented cytoDAPI with a custom Cellpose4 model (cell diameter = 15–23 pixels; flow threshold = 0.4; cell probability threshold = 0). 2P images are segmented in two dimensions from the mean of the motion-corrected recording, one image per functional plane, and the resulting masks are curated manually before registration. For the samples reported here, we use a custom Cellpose4 model (cell diameter = 7-9 pixels; flow threshold = 0.4, cell probability threshold = 0).

#### Low-to-high resolution 2P image registration

When a high-resolution image of the same 2P field of view used for functional imaging is supplied, EASI-PASS registers the functional 2P imaging data to this high-resolution image automatically. Matching features are found between the low and high-resolution 2P images by the scale-invariant feature transform (SIFT), and an affine transform is fit to them by random sample consensus (RANSAC). Residual nonlinear distortion within the field is then corrected with a dense optical-flow field. The functional image is now in the coordinates of the high-resolution 2P image, which is registered to the EASI-FISH volume by the cross-modal registration described below. Thus, applying these two transforms in sequence places the original functional masks into EASI-FISH coordinates. Masks are label images, in which each pixel value is an integer identifying one cell. Thus, to preserve cell identity, nearest-neighbor interpolation is used at every step.

### Cross-modal registration

#### Landmark initialization

Landmarks were placed between the 2P plane and the reference EASI-FISH volume using the ImageJ plugin BigWarp^9^. This is a critical step in the pipeline because erroneous landmark placement can limit registration quality in subsequent steps. A sparse red label is used to facilitate landmark placements, and typically 10-20 landmarks is sufficient to start the registration process. The landmarks are interpolated with a thin-plate spline that maps the flat imaging plane onto a smooth curved surface throughout the volume. Landmarks are placed by hand on a single 2P plane. When additional 2P planes are recorded at other depths, the same landmarks are used to initialize registration of each additional 2P plane. The 3D rigid body search described below shifts the hand-placed landmarks in depth and laterally, placing each additional 2P plane at its correct overall position in the EASI-FISH volume. Each additional 2P plane then proceeds independently through the remaining registration stages, which correct minor aberrations introduced by multiplane imaging with an electrically tunable lens.

#### 3D rigid body search

The rigid body search translates the 2P plane in x, y and z, scoring each offset by IoU between the 2P and EASI-FISH masks within the landmarked region (**Figure 2A**). Rotations are not included because they have already been captured by the thin-plate spline fit to the manual landmarks. Every combination of lateral and axial offset is evaluated. At each candidate depth the volume is resampled along the curved surface, and a cross-correlation in the Fourier domain returns the IoU at every in-plane offset in a single pass. The offset with the highest IoU over the whole three-dimensional grid is retained.

#### Global affine transform

For each 2P cell mask, we obtain a refined estimate of the corresponding EASI-FISH mask location. We then run an affine transformation using the matched masks as pairs of landmarks. To refine the estimates of each matched mask across the two datasets, a patch of 200 µm radius centered on each 2P cell mask is translated across the EASI-FISH volume (± 200 µm in X, ± 200 µm in Y, ± 75 µm in Z), and the IoU between the 2P and EASI-FISH masks is computed at each offset (**Figure 2B**). The offset that maximizes IoU is only retained if it exceeds the next-highest IoU in the window by 2%. An affine transform is fit to the retained displacements (i.e. the offsets between each pair of masks) by RANSAC (residual tolerance, 10 µm) and constrained to stretch or compress by no more than 20% in any direction. This transform maps the two dimensions of the 2P plane into the three dimensions of the EASI-FISH volume. Displacement in X, Y and Z varies linearly across the plane. This step corrects tilt, shear and scale error remaining after manual landmark placement.

#### Local refinement

The above steps for cross-modal mask matching and affine fitting and transformation are repeated sequentially for tiles of decreasing size (500, 200 and 100 µm by default). Thus, each iteration corrects for warping at a finer spatial scale than the last (see **Figure 2C**). For the first stage of the cascade we set the search window to 100 µm in X and Y and 30 µm in Z. Subsequent stages adapt the window to the residual misalignment measured at the previous stage. At every stage, the 2P field of view is divided into tiles overlapping by 50%. An affine transform is fit by least squares to the cell matches within each tile. Each fitted transform was required to stretch or compress the image by no more than 20% along any direction. A tile that exceeds the limit falls back to the median displacement of the cell matches in that tile. The resulting affine transform is accepted only if the tile meets all three of the following requirements: (1) the tile contains at least 3 matches, enough to fit the affine transformation; (2) applying the displacement does not reduce the tile’s IoU; and (3) the displacement in the tile does not differ from the median displacement of the eight nearest accepted tiles by more than four median absolute deviations. The median absolute deviation is calculated across the displacements of those eight tiles, as the median of the absolute differences between each of those displacements and the median displacement. Tiles that do not meet these conditions inherit the interpolated displacement field from the accepted tiles.

#### Building the displacement field

Tile displacements are interpolated to every pixel of the 2P plane with a thin-plate spline. Additional control points are placed along the plane perimeter at 100 µm spacing. Each control point is assigned a scaled version of the displacement of the nearest accepted tile, reduced linearly with distance from that accepted tile so that this perimeter control is not adjusted by an accepted tile spaced more than two tile widths away. Each pixel’s displacement is scaled down with distance from the nearest accepted tile. The scaling is a Gaussian with standard deviation σ = one tile width. Corrections are therefore confined to regions where tiles were accepted.

### Cell matching between 2P and EASI-FISH

Cells were matched between the 2P plane and the EASI-FISH volume by two independent methods, mask overlap and Soma-print, both of which are reported in the output table for every cell.

#### Mask overlap

After registration, the 2P plane is warped onto a curved surface within the EASI-FISH volume. Thus, a single 2P mask can intersect more than one EASI-FISH z plane. To compute IoU between a 2P mask, which is two-dimensional, and an EASI-FISH mask, which is three-dimensional, we counted only the part of the EASI-FISH mask lying within the planes that the 2P mask occupies. The rest of the EASI-FISH mask lies at depths the 2P plane never sampled. The intersection is the number of voxels the two masks share. The union is the size of the 2P mask plus the size of the restricted EASI-FISH mask minus the intersection. A neighborhood IoU is also reported for each 2P cell. All masks within a 50 µm square window centered on the cell are merged into one binary image in each modality, and the IoU between the two merged images is taken at the registered position, without searching over shifts. A high value indicates that the local arrangement of cells agrees between the two modalities.

#### Soma-print

Soma-print matches cells by the geometric arrangement of their neighbors^8^. Each cell is described by the two-dimensional vectors from its mask centroid to the centroid of its nearest neighbors; we used 15 neighbors for 2P cells and 30 for EASI-FISH cells, since cytoDAPI labels every cell and yields more *ex vivo* masks. To compare two cells, the Euclidean distance was computed between every pair of vectors drawn from the two sets, and the mean over the ten best-matching pairs is converted into a similarity score. Distant matches are penalized by multiplying that score by

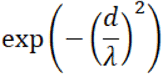

where d is the separation between the two centroids and λ is one tenth of the longer of the two 2P image dimensions. We compared a 2P cell only against EASI-FISH cells within 3λ (400 µm in most cases).

Because the 2P plane is transformed onto a curved surface within the EASI-FISH volume, the cells corresponding to it do not lie in a single flat plane. Scoring was repeated at three depths: the registered 2P surface, and the same surface displaced 5 µm up and down. Each cell was assigned the highest of the Soma-print scores across these three depths. As in the original method^8^, we used a likelihood-ratio threshold of 0.05 to select matches, so no absolute score cutoff is imposed (**Supplemental Figure 3A-B**). Matching is then iterated, with each cell’s neighborhood rebuilt using only cell masks that are confirmed to have matches in both datasets. Iteration stops when the number of matches changes by less than 5% from the previous iteration.

### Registration and matching across FISH rounds

We modified the computational workflow of the original EASI-FISH study^3^ for alignment across EASI-FISH rounds (outlined below). Alignment proceeds in two stages: a fast global stage that coarsely aligns the whole volume, and a slower local stage that corrects the deformation remaining within it. Both stages use cytoDAPI as a common stain present in every round.

#### Global registration

Each EASI-FISH round was registered to the reference EASI-FISH round using the centroids of the three-dimensional cell masks in both rounds, instead of blob-like image features^3^. Each cell is described by the cytoDAPI intensity pattern in a square window of 60 µm radius, centered on its centroid. We found this window size was large enough to be locally distinctive and small enough that this patch is unaffected by deformation elsewhere in the sample. Each patch is mean-subtracted and normalized, so the correlation between two patterns is their dot product. Every patch from cells in a subsequent round is correlated against every patch from the reference round. Each centroid of the later round is paired with the reference centroid of highest correlation, keeping only pairs with a correlation value of 0.5 or greater. A twelve-parameter affine transform was fit to the resulting centroid pairs by RANSAC with an inlier threshold of 15 µm. This stage runs on downsampled volumes and is therefore computationally inexpensive (<20 seconds per registration on a standard computer).

#### Selection

Because the global stage is fast, it was repeated over a small set of parameter settings, producing several candidate registrations for each round. Each candidate is scored visually by the user, as well as by the mutual information between the warped later round and the reference round, computed on the raw cytoDAPI images. Two additional metrics are reported per registration to help users assess registration quality: (1) number of centroid pairs that are mutually nearest neighbors within 10 µm and (2) the median distance between the centroids of those pairs. Candidates are ranked by mutual information. The user accepts the ranked choice or selects another, and this registration is used to initialize the local refinement step (see below).

#### Local refinement

A local 3D alignment stage then refined the EASI-FISH-to-EASI-FISH alignment block by block with a distributed piecewise registration. This stage starts from the selected global affine transform and runs at full resolution. The same two sets of cell centroids are reused, with the centroids of the later round mapped into the reference frame by the selected global transform. Three-dimensional blocks of 200 × 200 × 50 µm (200 × 200 × 10 voxels, with 50% overlap between neighboring blocks) were used throughout. We found that these parameters were a good starting point for all samples, though different block sizes can be specified for tissue whose cell density differs from that tested here. We did not find it necessary to include the final gradient-descent refinement step of the original EASI-FISH workflow^3^ and omitted it to reduce computational cost.

#### Matching

Cells were matched between the reference round and a later round in two stages. Each cell of the later round was first scored against reference cells within a local window by best single-plane IoU. Cell pairs that were each other’s best match, with IoU above 0.5, were accepted as seed matches. These seed matches were held fixed while Soma-print ran on the cells left unmatched, evaluating candidate matches for each cell from neighbors within ±2 planes, or 10 to 15 µm.

### mRNA fluorescence extraction and output

For each EASI-FISH round and color channel, we extract the mean intensity of the pixels in each cell’s mask, as well as an estimate of the mean local background intensity. Each cell’s background intensity estimate is extracted from a square window of 30 µm radius in X and Y, centered on the cell’s centroid in each plane the cell occupies. Pixels belonging to any cell mask are excluded, along with a one-pixel perimeter around each mask. The resulting data are combined into a table with one row per cell of the reference EASI-FISH round. Columns indicate the per-cell and neighborhood IoU of the 2P partner with the highest overlap, the matched 2P cell identities for Soma-print and IoU-based matching, Soma-print outputs (score and confidence), and mask dimensions.

## QUANTIFICATION AND STATISTICAL ANALYSIS

### Registration and matching quality

Alignment quality metrics (**Figure 3** and **Supplemental Figures 2-4**) were evaluated at three registration stages: (1) after landmark-based warping (which we refer to as baseline), (2) after the 3D rigid body search followed by the global affine transform, and (3) after the subsequent stages of local non-rigid refinement. For additional planes other than the initially landmarked plane, the 3D rigid body search was used as the baseline to correct for the different depth of each functional plane.

### Analysis of slice imaging experiments

#### Inclusion criteria

Analyses were restricted to a manually drawn boundary around the PBN in each slice, thereby excluding neurons near the edge of the imaging field and, in the EASI-FISH volume, neurons near the edge of the tissue.

#### Response classification

For each cell, the mean ΔF/F over the 5 s following stimulation was compared with the mean over the preceding 10-s baseline across trials by a paired t test. Neurons were classified as significantly activated at p < 0.05.

#### Gene expression

Gene expression was measured for each probe as the mean fluorescence within each cell in that probe’s channel, minus the local background (see mRNA fluorescence extraction and output). Negative values were set to zero. Intensities were then centered on the median across cells and scaled by the spread of the middle half of cells. A two-component Gaussian mixture was fit per gene and per animal across the EASI-FISH volume, and the threshold was applied within the region of interest. Cells assigned to the higher component were identified as positive for that gene.

#### Expression gradients

Within each animal, neurons were ranked by their expression of a given gene, and binned into deciles (i.e. into ten bins with the same number of cells per bin) according to expression level. The fraction of neurons classified as significantly activated (paired t test, p < 0.05) was computed per bin and averaged across animals with equal weight per animal. Stimulated responses were averaged across cells within each bin. The population mean trace of the same animal and gene was then subtracted, giving an adjusted trace. Traces were smoothed with a 1-second moving average window (**Figure 4K-M**).

#### Conditional probabilities

For each gene, the fraction of significantly activated neurons that were gene-positive and the fraction of gene-positive neurons that were significantly activated were computed per animal using the binary classification explained above. These fractions were then averaged across mice (**Figure 4N**).

#### Regression against neuron location and ChrimsonR axon density

A linear mixed effects model was fit for each gene across all neurons to test whether gene expression predicted a cell’s evoked response beyond its position within the tissue and its proximity to labeled afferents. The dependent variable was the evoked ΔF/F, z-scored within animal. Predictors were expression of that gene, also z-scored within animal; the cell’s position within the tissue; and the local ChrimsonR-tdTomato signal. Position was entered as a quadratic surface, comprising both coordinates, their squares and their product, which absorbs any smooth spatial gradient of excitation. The ChrimsonR-tdTomato signal was taken as the sum of the mean intensity within the cell and the mean intensity within the surrounding background region. Animal identity was entered as a random intercept, with a random slope for gene expression. Each gene was tested separately, p values were corrected across genes by the Benjamini-Hochberg procedure, and effects are reported as the change in z-scored evoked ΔF/F per standard deviation of expression (**Figure 4O**).

#### Comparison with neighboring neurons

Within each sample, and for each gene, high-expressing neurons were defined as those in the top decile of expression and low-expressing neurons were defined as those below the median expression. Each high-expressing neuron was paired with its five nearest low-expressing neurons (median neighbor distance 45.3 µm). For each of these neurons we computed the neighborhood change, defined as the neuron’s evoked response (ΔF/F) minus the mean evoked response of its five neighbors (**Figure 4P**). A gene and sample were analyzed only if at least 25 high-expressing neurons had low-expressing neighbors within a median distance of 50 µm. Neurons whose nearest low-expressing neighbors lie farther away do not provide a local comparison. Neighborhood changes in evoked ΔF/F were tested with a linear mixed-effects model with sample as a random effect, and p values were corrected across genes by the Benjamini-Hochberg procedure.

#### Statistical testing

Odds ratios for the association between gene expression and activation were computed within each mouse and then combined, so that differences in baseline activation between mice do not drive the pooled estimate (Mantel-Haenszel method). Across genes, p values were corrected by the Benjamini-Hochberg procedure.

### Analysis of *in vivo* cortical imaging data

#### Cell-type assignment

Each EASI-FISH cell was matched to the 2P cell with which it had the highest intersection over union, restricted to the 2P plane the cell occupies (see above). For a given gene marker in EASI-FISH data from cortex (*Pvalb*, *Sst*, *Vip* and *Calb2*), we estimated the mean intensity of that marker across all pixels belonging to a given cell. The mean intensity of the background surrounding the cell was then subtracted and negative values were set to zero. Because staining intensity varied between animals and probes, cells positive for a given gene were identified by manually defining a threshold that divided bimodal peaks in the log intensity of HCR signal.

Class assignment was exclusive and ordered: cells were first chosen to be SST+ or SST-, then VIP neurons were selected among those neurons not already assigned as being SST+, then PV neurons were selected among those neurons not already assigned as being SST+ or VIP+. All other matched neurons were defined as unlabeled (putative excitatory). The presence or absence of *Calb2* expression was independently assessed and used to subdivide the SST and VIP classes.

#### Pupillometry

Keypoints tracked with confidence below 0.90 were discarded. Gaps of up to 14 frames (approximately 0.5 s) were filled with a duplicate of the last high-confidence frame. Longer gaps were left as missing. An ellipse was fit to the pupil keypoints in each frame by constrained least squares. Pupil area was taken as πab, where a and b are the semi-axes of the fitted ellipse. Frames whose area deviated from the session mean by more than 4 SD in either direction were removed and filled by linear interpolation. Eyelid width was measured once per session, as the distance between the eyelid keypoints of the median pose across all frames. Frames in which the fitted pupil area exceeded half the area of a circle with a diameter equal to the eyelid width were removed as blinks. The trace was scaled to minimum and maximum values of 0 and 1, then averaged over the three frames acquired during each imaging volume.

#### Arousal coupling

We measured coupling of neural activity to pupil-linked arousal as follows: For each cell, we estimated the Pearson correlation between the smoothed deconvolved activity of a neuron and zero-lag pupil area across all “spontaneous” frames, defined as all frames preceding the first visual stimulus together with all frames from 10 s after the last stimulus until the end of the recording.

#### Spatial correlation decay

Pairwise Pearson correlations between neurons of the same class were computed over the same set of “spontaneous” frames as described above. To remove cell-cell variability in distance-independent correlation, each cell’s correlations were normalized by subtracting the mean of that cell’s correlations with all other cells, regardless of class. Pairs used for fitting were then restricted to cells of the same class. Distances were estimated as lateral Euclidean distances between mask centroids in micrometers, pooled across imaging planes, with differences in depth excluded from the metric. For each cell with at least 10 within-class partners, the within-class activity correlation was fit by least squares as a function of lateral separation, d, as

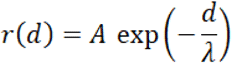

where A is the pairwise correlation at zero separation and λ is the decay constant. Fits were bound to decay constants of 5 µm to 1,000 µm, and fits converging at a bound were discarded. Putative excitatory neurons were randomly subsampled to 1,000 cells per animal before fitting. Within-class decay constants from individual cells were pooled across animals within each class and summarized per animal as the median for per-animal tests and as the geometric mean for plotted summaries (**Figure 5I-J**). Qualitatively similar results were obtained with and without a distance-independent intercept term in exponential fit equation (data not shown). Because fit intercepts were small (on the order of 0.01), we used the version of the equation without the intercept term for simplicity.

#### Control for fit quality

We considered whether classes might systematically differ in how well the falloff in within-class correlation with distance was fit by an exponential function, which would bias comparisons of the decay constant. Fits were therefore repeated with the amplitude restricted to values between 0 and 1. Neurons were also resampled from each class so that all classes matched a common distribution of goodness of fit. Main analyses used all fits that converged and did not hit any parameter bounds, without resampling.

#### Statistical testing

Differences across classes at the level of individual neurons were evaluated using a Kruskal-Wallis statistic against a null generated by permuting class labels within animal (1,000 permutations), which prevents differences between animals from driving the result. Putative excitatory neurons were excluded from this test. Differences at the level of summary statistics were evaluated using Friedman tests (**Figure 5F-G, I-J**). CALB2 subtype comparisons were tested using within-animal differences between *Calb2*+ and *Calb2-* neurons, against a null generated involving permutation of subtype labels within animal.

#### Sample sizes

All five animals contributed to the pooled distributions computed across neurons. Per-animal tests use four animals: one animal lacked two of the EASI-FISH probes and was excluded from analyses requiring all probes. For **Figure 5J**, we excluded one mouse lacking sufficient subtype neuron counts to meaningfully assess differences in exponential fits.

All data are reported as mean ± SEM unless stated otherwise.

---

**Supplemental Figure 1.**
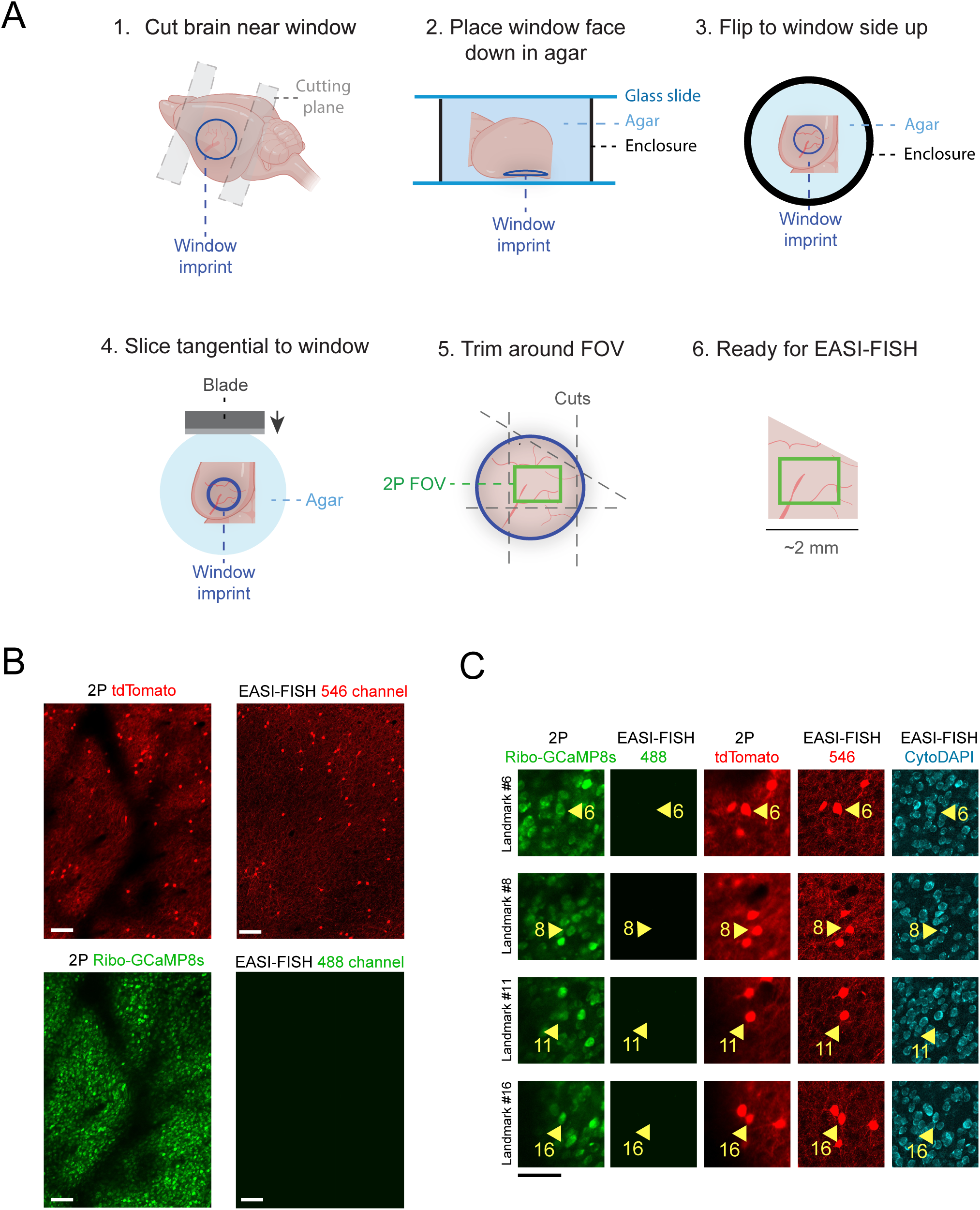
Additional EASI-FISH tissue processing details. Related to Figures 1 and 2. **(A)** Extraction of the imaged cortical volume. After the final imaging session, animals were perfused and the brain was extracted. A large tissue block (3 × 3 × 0.25 mm) was cut tangential to the imaged region, using the cranial window imprint as the physical reference for cutting tangential to the cortical surface. This block was then trimmed under a stereoscope, and blood vessels were used to guide asymmetric cuts to retain the correct orientation during EASI-FISH. **(B)** Example field of view from an *in vivo* experiment, showing preservation of tdTomato and loss of Ribo-GCaMP8s fluorescence after the first round of EASI-FISH. Left panels: *in vivo* two-photon (2P) images. Right panels: *ex vivo* EASI-FISH images. **(C)** Four example landmarks from (B), identified using tdTomato. Columns show 2P Ribo-GCaMP8s, EASI-FISH 488, 2P tdTomato, EASI-FISH 546 and EASI-FISH cytoDAPI. Yellow arrowheads mark each landmark. Scale bars, 100 µm unless otherwise noted.

**Supplemental Figure 2.**
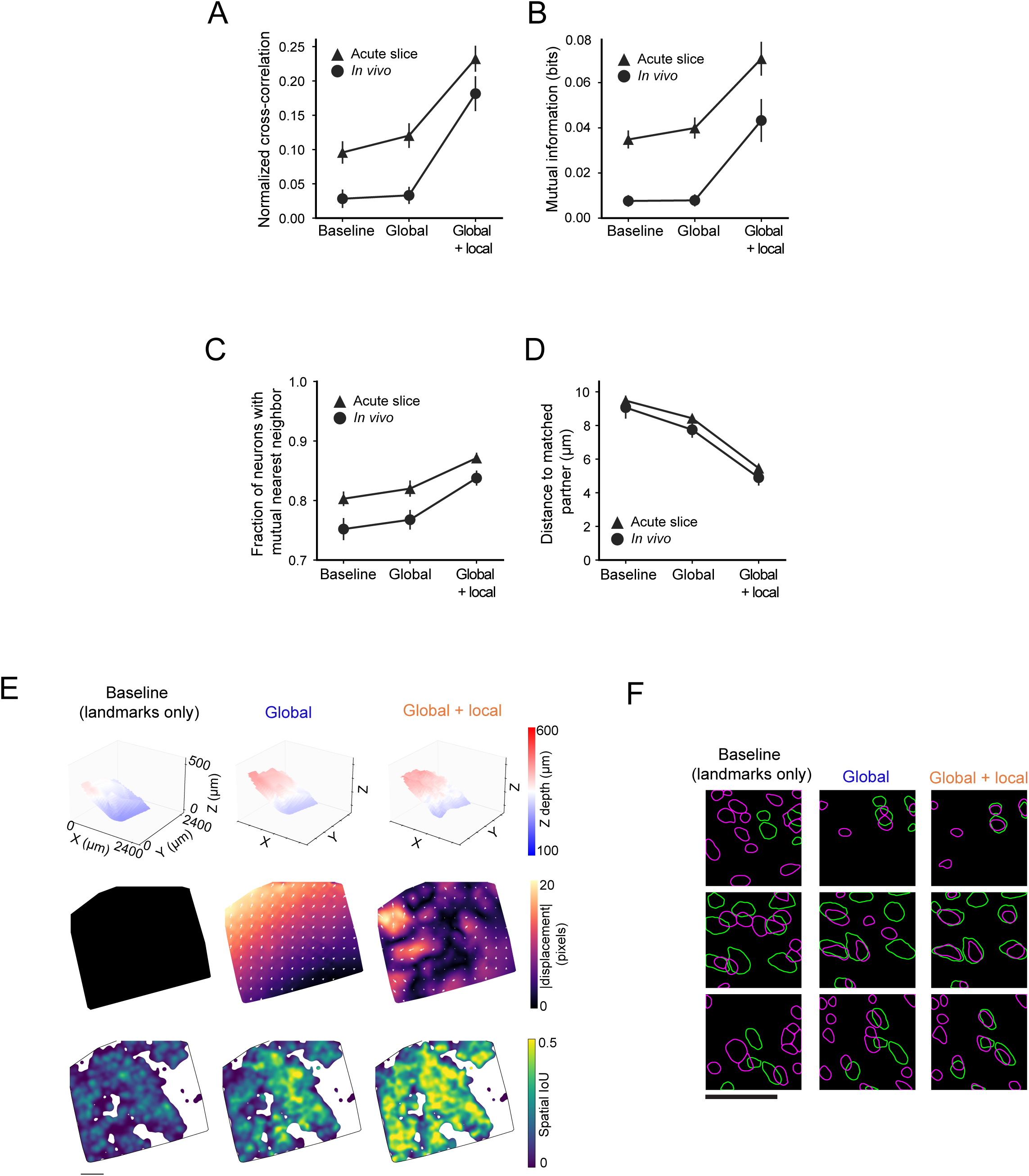
Additional registration performance metrics for image alignment from 2P to EASI-FISH. Related to Figures 2 and 3. **(A)** Normalized cross-correlation between 2P images (Ribo-GCaMP8s) and corresponding EASI-FISH surface (cytoDAPI). **(B)** Mutual information between 2P images (Ribo-GCaMP8s) and corresponding EASI-FISH surface (cytoDAPI). (A) and (B) were computed from image intensities rather than from segmented masks. **(C)** Fraction of putatively matched neurons across the two modalities that are mutual nearest neighbors, meaning each cell of a pair is the closest partner of the other cell. **(D)** Distance between the centroids of putatively matched cells. For (A) to (D), triangles, acute slice imaging (n = 7 slices from 7 mice). Circles, in vivo imaging (n = 5 mice). **(E)** Example 2P field of view across registration stages (baseline, global, global and local). Baseline indicates initial alignment using manual landmarks. Top row: 3D rendering of the warping of the 2P plane within the EASI-FISH volume. Middle row: displacement field at each stage. Bottom row: spatially smoothed IoU between 2P and EASI-FISH masks. **(F)** 2P (green) and EASI-FISH (purple) mask contours for 3 example regions across alignment stages. Data are mean ± SEM across samples. Error bars in (A) to (D) show SEM. Scale bars, 100 µm.

**Supplemental Figure 3.**
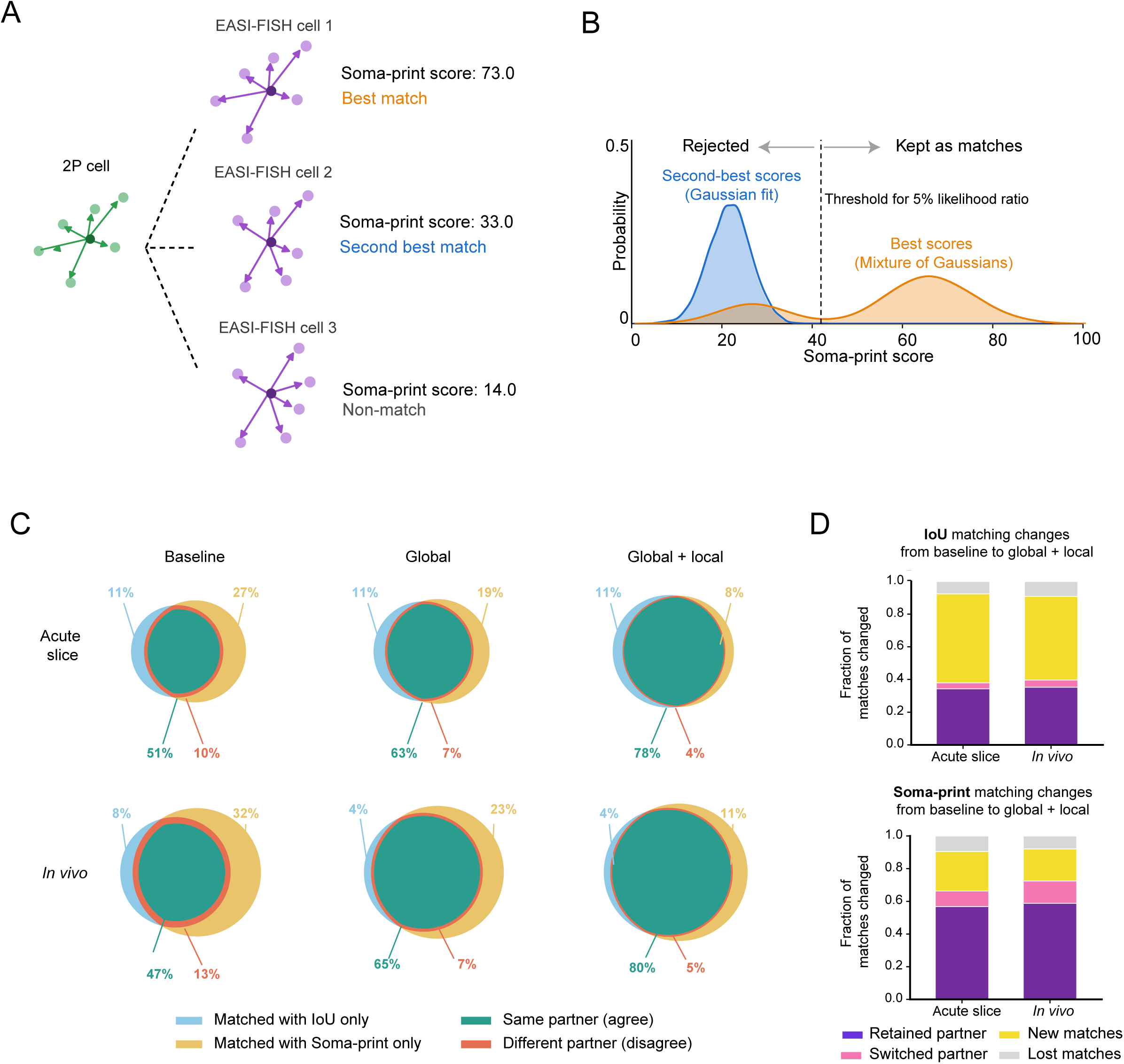
Matching performance based on IoU or Soma-print across EASI-PASS registration stages. Related to Figure 3. **(A)** Explanation of the Soma-print algorithm. Each cell is described by the vectors from the centroid of that cell to the centroids of the nearest neighbors of that cell. A 2P cell is scored against every candidate EASI-FISH cell by the similarity of the two sets of vectors. The highest-scoring candidate is the best match and the next-highest is the second-best match. **(B)** Selection of confident matches. For every 2P cell, the best and second-best Soma-print scores are recorded. The distribution of second-best scores is fit with a single Gaussian (blue), which serves as the null distribution for scores arising by chance. The distribution of best scores is fit with a two-component Gaussian mixture (orange). For each cell, the likelihood ratio is the density of the best score of that cell under the null distribution divided by the density under the mixture. A cell is kept as a match when the likelihood ratio falls below 0.05 (dashed line). **(C)** Fraction of cells for which both Soma-print and IoU matched the same EASI-FISH partner, across EASI-PASS registration stages. Acute slice, n = 20,397 neurons; *in vivo*, n = 27,152 neurons. **(D)** Change in the fraction of cells matched by IoU and by Soma-print, from baseline to global and local alignment. Acute slice, n = 20,397 neurons; *in vivo*, n = 27,152 neurons.

**Supplemental Figure 4.**
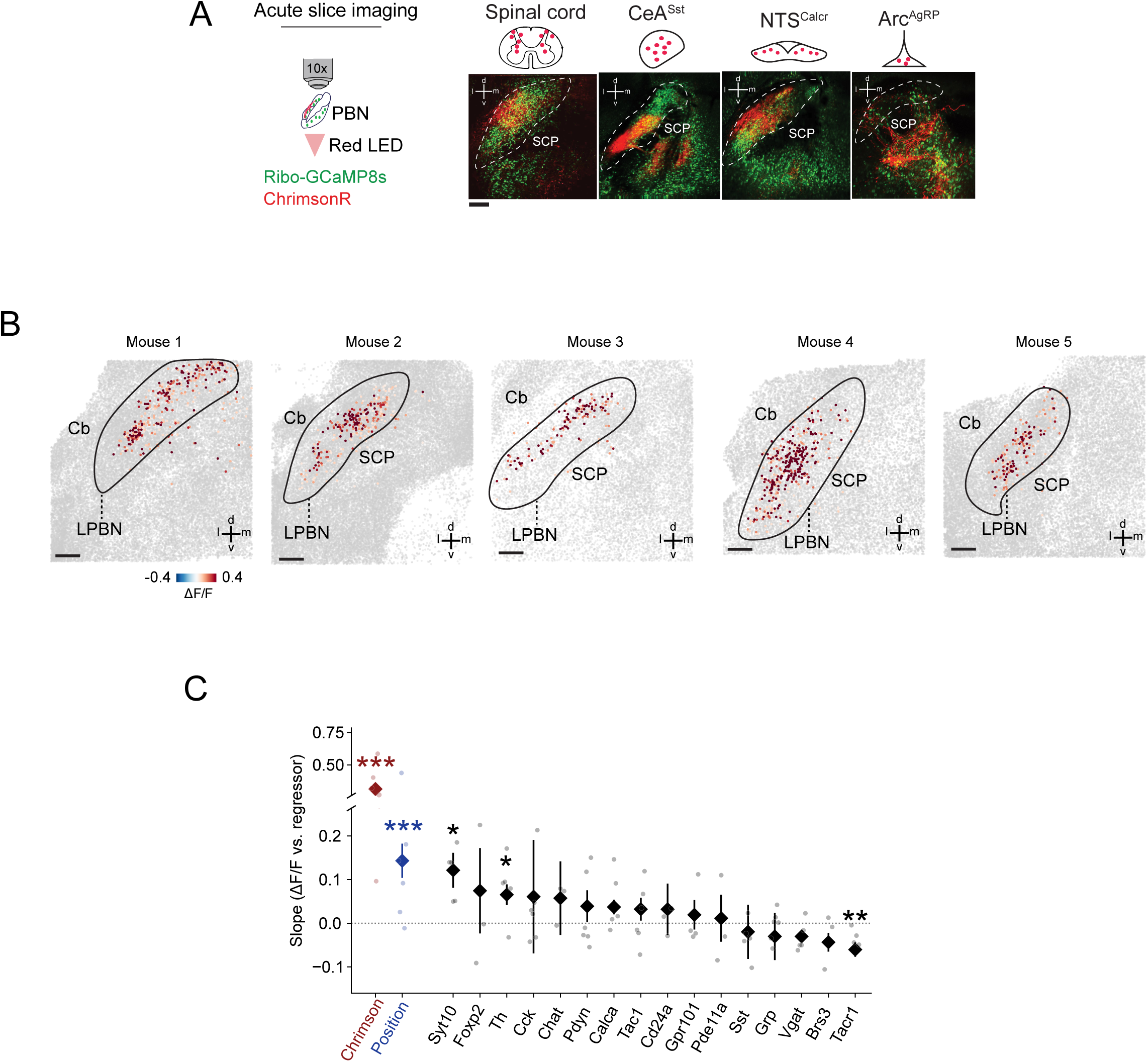
Supplemental analyses of acute slice imaging experiments in the parabrachial nucleus. Related to Figure 4. **(A)** Example acute slice imaging fields of view from experiments where ChrimsonR was expressed in different input populations, leading to expression of tdTomato in long-range axons in PBN. Ribo-GCaMP8s is also expressed in all neuronal cell bodies. Input populations shown are lumbar spinal cord, CeA-Sst, NTS-Calcr and Arc-AgRP. **(B)** Centroids of PBN neurons significantly activated by stimulation of spinal inputs for example slices from n = 5 mice, overlaid on a maximum projection of all EASI-FISH cells (gray) after registration. Centroids are colored by evoked ΔF/F. Cb, cerebellum; SCP, superior cerebellar peduncle. **(C)** Regression slope of evoked ΔF/F on gene expression, soma position and local ChrimsonR expression. Linear mixed-effects model over all matched cells (n = 9,024 neurons from 6 mice), with mouse as a random effect. Note the broken y axis. Diamonds, fixed-effect estimate; error bars, one standard error of that estimate. Gray dots, per-mouse slope. Red, local ChrimsonR expression; blue, soma position; black, gene expression. Stars, significance after Benjamini-Hochberg correction within regressor family: *q < 0.05, **q < 0.01, ***q < 0.001. Data are mean ± SEM across mice unless otherwise noted. Scale bars, 100 µm.

**Supplemental Figure 5.**
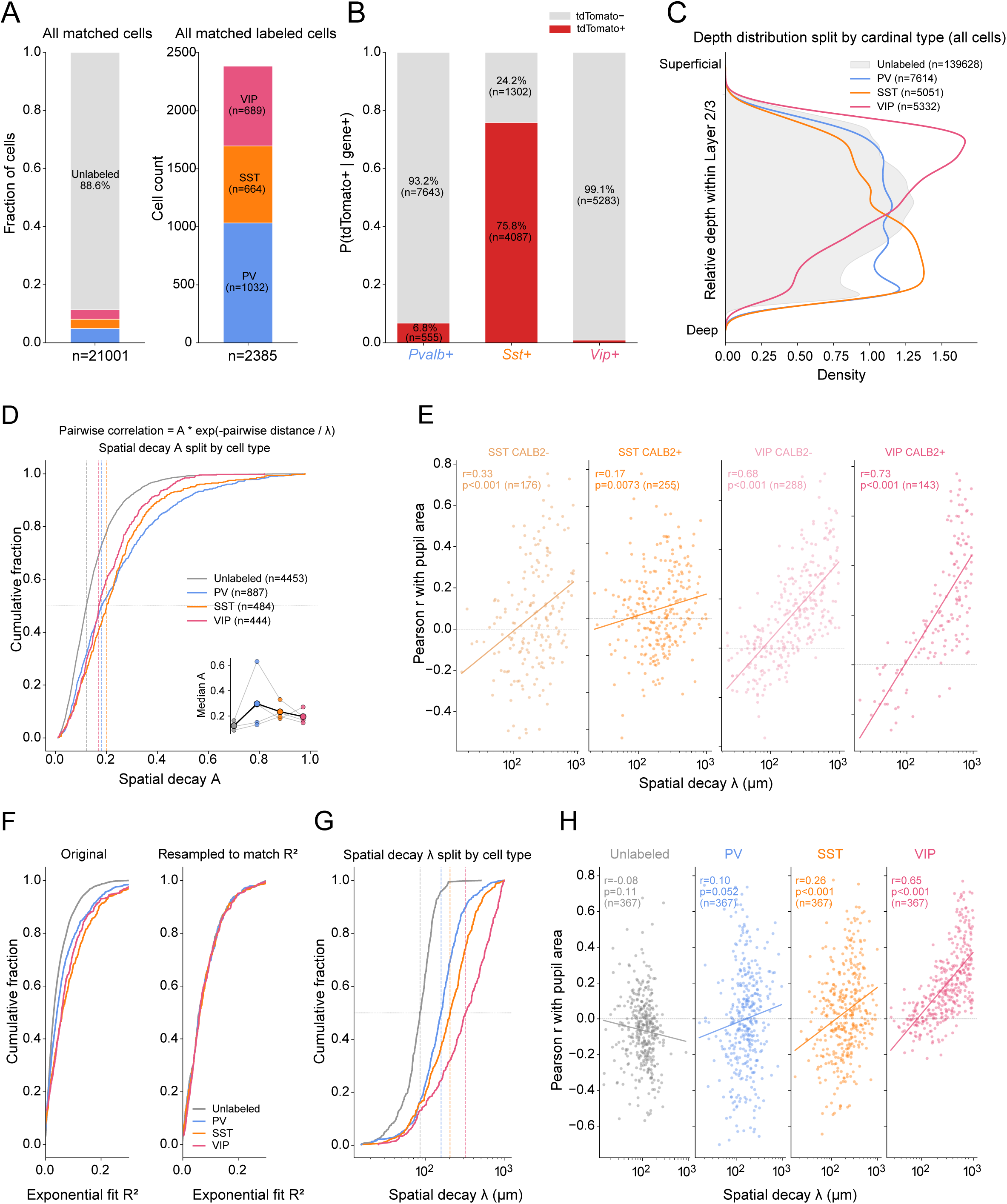
Further analysis of EASI-PASS applied to the visual cortex. Related to Figure 5. **(A)** Left: pooled fractions of cells belonging to each cell class, among all matched cells. Right: pooled counts of all labeled cells, split by cardinal class (n = 5 mice). **(B)** Fraction of tdTomato-expressing cells, split by cells with EASI-FISH expression of *Pvalb*, *Sst*, or *Vip*. (B) and (C) use the full EASI-FISH tissue, with no restriction to matched cells. Only mice with all cardinal classes represented are included (n = 4 mice). **(C)** Distribution of relative depth within layer 2/3 for each cardinal class. Depth differed across classes: Kruskal-Wallis within-mouse permutation test, H = 1147.03, p < 0.001, n = 17,997 neurons from 4 mice. **(D)** Distribution of the fitted amplitude A of the spatial decay exponential fit (i.e. the pairwise correlation extrapolated to zero inter-pair distance), split by cardinal class. Kruskal-Wallis within-mouse permutation test, H = 5.16, p = 0.082, n = 1,815 neurons. Friedman test on per-mouse medians across classes, χ² = 7.50, p = 0.058, n = 4 mice. **(E)** Correlation between λ and pupil-area correlation, for *Calb2*-defined subtypes of SST and VIP neurons (n = 4 mice). **(F)** Resampling approach used to match R² across cardinal classes, where R² is the goodness of the exponential decay fit for each cell (see STAR Methods). **(G)** λ split by cardinal class, after resampling to match R². **(H)** Within-class correlation between λ and pupil modulation, after resampling to match R² (n = 4 mice). In the inset to (D), gray lines are individual mice and the black line is the mean across mice.

